# Factorization of Alzheimer’s disease genetic risk influences allow patient stratification, predicting disease onset, cognitive decline, and cell-type specific responses

**DOI:** 10.64898/2026.01.02.697432

**Authors:** Alexandre Pelletier, Tony Tuck, Julia TCW

## Abstract

Late-onset Alzheimer’s disease (AD) is a complex and heterogeneous neurodegenerative disease with significant genetic components implicated in at least 97 loci from AD genome-wide association studies. While various distinct AD subtypes have been identified based on brain or CSF molecular profiling, contribution of the genetic signatures in distinguishing the AD subtypes is lacking. Here, we leveraged large snRNA-seq postmortem brain data with an empirical Bayes matrix factorization (EBMF) approach to study common effects of 197 AD risk variants on neuronal and glial cell transcriptome, enabling factor-based polygenic score (fPGS) and patient clustering based on their functional genetic profiles. We confirmed that each factor captures specific AD risk variant influences on cell types and known AD-associated biological processes, such as mitochondrial activity, endo-lysosomal activity, mRNA processing, neuroinflammation, or calcium signaling. Further, we found that most fPGS were predicting a certain neuropathological or AD-associated molecular condition. Notably, fPGS3 predicts somatic mutation burden in excitatory neurons, and fPGS7 predicts epigenome erosion in excitatory neurons associated with lipid transport disorders, increased mitochondrial activity, and increased Tau pathology. Finally, unsupervised clustering analysis of individuals with mild cognitive impairment and AD based on their fPGS profiles enable us to identify seven clusters, which are differentiated by the *APOE* genotype. Among them, five groups were subdivided within *APOE3* homozygotes, which remarkably predicted either the disease severity or the disease onset with up to 6 years differences among groups. Resilient groups were associated with reduced matrisome and reactivity in astrocytes and increased cholesterol metabolic pathways in oligodendrocytes and OPCs. Overall, the analyses confirmed the ability of EBMF to stratify AD risk variant influences into molecularly and clinically coherent factors that allow genetic predictions of particular cell types and alterations of biological processes impacting the disease.

## Main

Late-onset Alzheimer’s Disease (AD) is a complex neurodegenerative disease involving several molecular pathways and vicious feedback loops between neurons and glial cells along the progression of the disease. Even if AD is characterized by neurotoxic oligomeric amyloid-β and Tau tangles propagating in the brain, the molecular processes initiating this process are complex and heterogeneous across individuals. Mitochondrial dysfunction, chronic neuroinflammation, calcium signaling impairment, endo-lysosomal system breakdown, cholesterol/lipid dysregulation, or blood brain barrier leakage are all demonstrated mechanisms contributing to the disease^1–6^, however, the relative contribution of each processes appear to be divergent across various AD models or patient populations^7,8^. Recent studies have found that AD patients can be grouped into different subtypes based on their proteomic profiles affected in cerebrospinal fluid (CSF)^9^, brain^8^, or plasma^10^ that are associated with various clinical outcomes, further supporting the importance of the disease heterogeneity.

A significant part of the AD onset variability in the aging population is explained by genetics, as demonstrated by twin studies evaluating two-thirds of the genetic heritability of the disease^11^. To date, at least 97 loci have been found across more than 17 AD genome-wide association studies (GWAS), implying genes involved in diverse cellular processes, likely capturing a representative spectrum of the molecular drivers of the disease. Polygenic scores (PGS) combining the genetic risk of each individual have shown strong AD onset predictability^12^ but lack of molecular insights. Previous work generating cell-type specific PGSs have meanwhile found distinct astrocytic and microglial influence on AD endophenotypes^13^, however limited to a subset of the risk loci, notably excluding *APOE*, the leading AD risk locus, which limits a discovery of resilience factors or multicellular and coordinated effects. Here, we hypothesize that all AD risk loci can be grouped according to their diverse impact on AD pathology by unbiased assessments of their co-influence on brain cell types using postmortem single nucleus transcriptome data. Using an Empirical Bayes Matrix Factorization (EBMF) approach weighting influences of single nucleotide polymorphisms (SNPs or variants) in AD risk loci on functionally relevant factors, we found that groups of SNPs associate with distinct cellular and molecular pathways and AD pathologies, allowing us to stratify AD patients into clinically coherent groups. Notably, we identified five functional genetic clusters within *APOE* 33 individuals subdivided by specific molecular signatures, which differentiate disease onset and severity, leading to discovery of a disease resilience group and its cellular functions.

## Results

### EBMF factor-based AD SNP clustering predicts cell-type and functional influences

To functionally characterize the different genetic risk of AD and stratify AD patients accordingly, we conducted a data-driven study of the common influence of AD risk SNPs on brain transcriptomes (Figure 1). First, we assessed the impact of each AD risk allele on the brain cell-type level transcriptome using differential expression analysis (1). The resulting z-score matrix of the variant effects on each cell-type-gene (CtG) is factorized into two low-rank matrices: the factors (F) and their associated gene loadings (L) using an Empirical Bayes Matrix Factorization (EBMF) method^19,21^ (2). We then performed functional enrichment analyses by over-representation analysis (ORA) or fast-rank gene set enrichment analysis (fGSEA) on gene loadings (L) to characterize cell types and molecular pathways captured by each factor (3). We used the SNP weights/contribution for each factor to generate functionally coherent factor-based polygenic scores (fPGSs) and characterized these fPGSs for their association with clinical and molecular AD and aging traits including clinical traits, cell-type proportion, epigenome integrity, and DNA damage burden (4). Finally, we used the fPGS profile of each individual with mild cognitive impairment (MCI) or AD to cluster them and identify AD-relevant genetic subgroups that are risk or resilient for AD (5).

**Figure 1.**
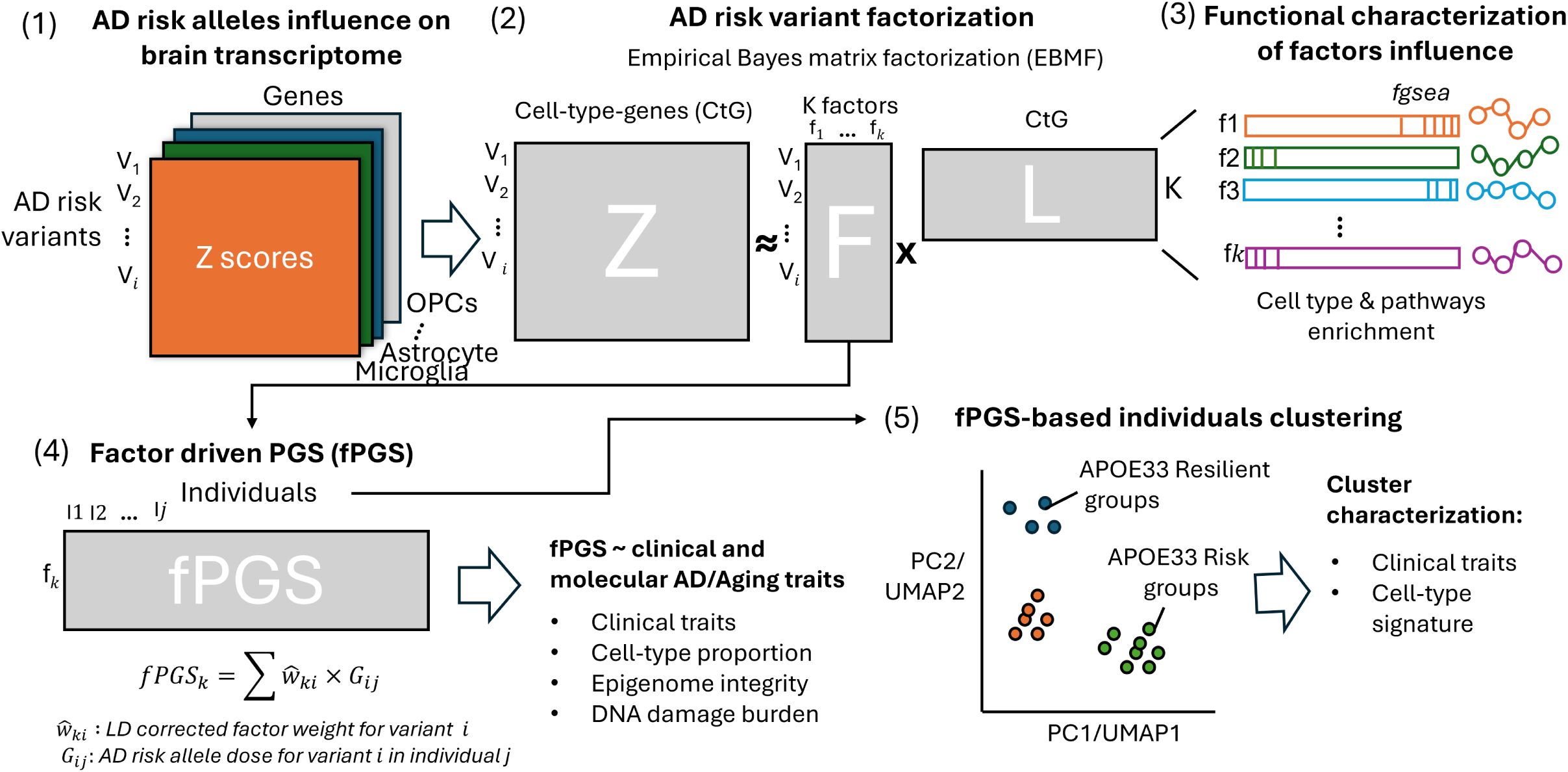
Overview of the analytical workflow. We evaluated how each AD risk allele affects brain cell-type transcriptomes via differential expression analysis, producing a z-score matrix of variant effects per cell-type-gene (CtG) (1). Using Empirical Bayes Matrix Factorization (EBMF), we decomposed this matrix into two low-rank matrices: factors (F) and gene loadings (L) (2). We then characterized each factor’s biological relevance through functional enrichment analyses (ORA or fGSEA) on the gene loadings (3). Next, we derived factor-based polygenic scores (fPGSs) from SNP weights, linking these scores to clinical and molecular AD/aging traits (4). Finally, we clustered patient based on their fPGS profiles to identify genetic subgroups associated with AD risk or resilience.

To perform this study, we used prefrontal cortex (PFC) of postmortem brain snRNA-seq data and matching whole genome sequencing from 372 aged individuals of the Religious Orders Study/Memory and Aging Project (ROSMAP) Mathys *et al.* study^14^. We used a representative panel of 237 leading SNPs of AD risk loci found across 26 AD GWAS available in the GWAS catalog (Supplementary Table 1) and tested their influences on 18 brain cell types and cortical layers, using a common cell-type definition by the BRAIN Initiative Cell Atlas Network (BICAN)^15^. We performed the differential expression analysis per cell type for all SNPs testing the effect of the AD risk allele dosage using DESeq2 (adjusting for the number of cells, mitochondrial percentage, sex, age-at-death, PMI, and ancestry). Following QC to ensure the removal of variants with strong LD (R^2^ > 0.5) without distinct transcriptome influence, 173 variants including 24 variants (14%) present in the *APOE* locus were preserved (Supplementary Figures 1A, Supplementary Table 2). In order to capture the putative haplotype effect, we intentionally preserved the SNPs that were in moderate LD with others (inclusion criteria: R^2^_LD_ < 0.5 or R^2^_zscore_ < 0.5 or delta between R^2^_zscore_ and R^2^_LD_ > 0.1; Supplementary Figure 1B, see Methods). We found distinct patterns of differentially expressed genes (DEGs; per SNP FDR < 0.05 and |log2FC| > 0.25) across cell types, with higher numbers of DEGs found in Exc L2/3 IT and Exc L5 IT, followed by astrocytes (Supplementary Figure 1C, Supplementary Table 3). Interestingly, the variant with the broader impact on brain transcriptome was the TMEM106B intronic variant rs13237518 and was predominantly impacting expressions on neuron subtypes (Supplementary Figure 1C).

Taking the entire SNP matrix of transcriptome influence (z-score), we used the EBMF approach to find common patterns of SNP influence across genes and cell types. We first assessed the quality of the method in capturing functionally relevant patterns among SNPs by performing cluster analyses. We used the first six factors (f1-f6; for example, f1 explaining for ∼ 8% and f2 for ∼ 6% of the total variance) (Supplementary Figure 1D) and clustered the SNPs based on the six factors using a Share Nearest Neighbor (SNN) graph and a leiden algorithm at two resolutions. Compared with a standard principal component analysis (PCA) using the first six principal components or a K-means (k = 100) followed by hierarchical clustering, we found that the first six factor-based EBMF was able to identify significantly more SNP clusters with greater silhouette scores relative to PCA or K-means clustering (Figure 2A). Further, each cluster SNPs were associated more strongly with specific cell types or biological processes after performing functional enrichment on the top 2,000 most specifically influenced genes per cluster (Figures 2B-2C). Using the higher clustering resolution leiden1.3 rather than leiden1, we identified eight clusters of SNPs with each one preferentially affecting certain cell types (Figures 2D-2E, Supplementary Table 4). Notably, cluster 1 was mostly associated with excitatory neuron L2/3 and L5 IT and astrocyte; cluster 2 with excitatory neurons L6 IT and L6b, oligodendrocyte, oligodendrocyte progenitor (OPC), and astrocyte; cluster 3 with inhibitory neurons PvalB and Vip and excitatory neuron L2/3; cluster 4 with inhibitory neurons Lamp5 and Sst and excitatory neuron L2/3 IT; cluster 5 with oligodendrocyte, OPC, and microglia; cluster 6 with oligodendrocyte, microglia, and inhibitory neuron PvalB; cluster 7 with several inhibitory neurons and microglia; and cluster 8 with all layer 6 excitatory neurons, oligodendrocyte, and OPC (Figure 2E). Concordantly, we found that cluster 1 (including SNPs in the *HS3ST1*, *TNIP1*, *NME8*, *FERMT2*, and *SORL1* loci) was regulating genes associated with neuronal activity and synaptic regulation; cluster 2 (*BIN1, INPPD, CLU,* and *APP*) with autophagy and mRNA catabolism; cluster 3 (*ACE*, *CR1*, *MAF*, *MS4A4A*, and *CTSB*) with mitochondrial activity, lysosomal acidification, and mRNA splicing; cluster 4 (*ABCA7*, *PTK2B*, and *CTSH*) with protein catabolic process, oxidative metabolism, and response to stress; cluster 5 (*TREM2, CD33, HLA-DRB1/DQA1, HS3ST5,* and *MEF2C*) with immune cell activation, response to lipid, endo-lysosomal pathways, and Tau protein kinase activity; cluster 6 (*CHRNE*, *ABCA1*) with cholesterol biosynthesis; cluster 7 (*PLCG2*, *GRN,* and *IL34*) with response to morphine and retinol binding; and cluster 8 (*FAM47E* and *CASS4*) with ribosomal activity (Figures 2D and 2F, Supplementary Table 5). Interestingly, some variants present in the same locus (e.g. *BIN1* or *APOE*) were not necessarily clustered in the same group (Figure 2D). For example, BIN1|NIFKP9 (rs6431220) and BIN1 (rs744373) was belonging to synapse-associated cluster 1 and autophagy-associated cluster 2, respectively (Figures 2D and 2F), suggesting those variants could have a distinct effect on AD while being genomically in a close proximity.

**Figure 2.**
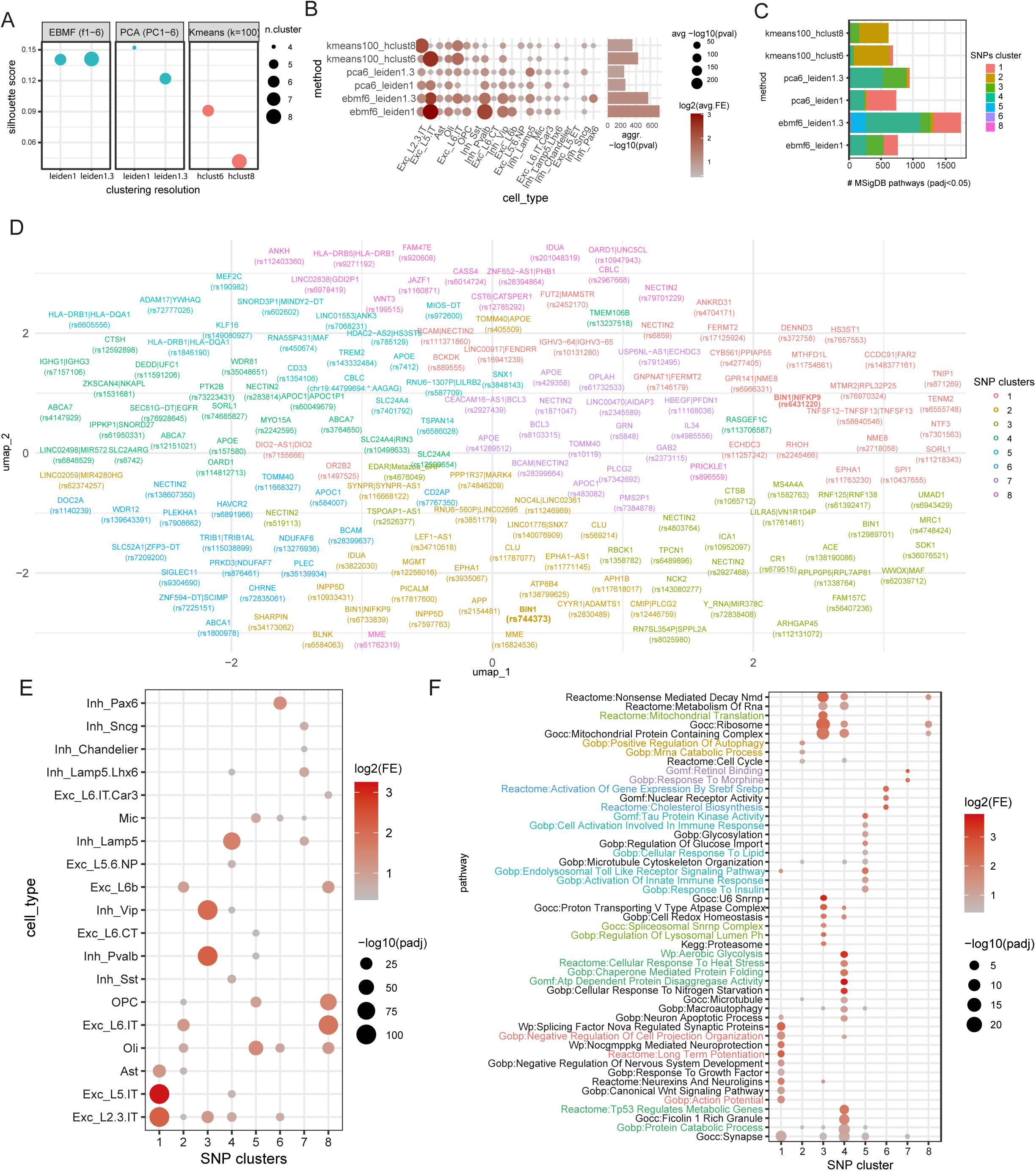
EBMF-based factorization and functionally relevant AD SNP clustering based on common impacts on brain cell-type transcriptome. **A**: SNP clustering performance comparison based on EBMF, PCA, or Kmeans. For EBMF and PCA, network clustering is based on the first six dimensions (i.e. factors 1 to 6 and PCs 1 to 6) using the leiden algorithm at resolution 1 or 1.3. For K-means, generating 6 or 8 clusters using hierarchical clustering (hcluster) on k = 100 centers. **B-C**: For each method, the average cell-type enrichment (C) and number of MSigDB pathways (GO and CP) enriched in the top 2,000 cell-type-genes (CtG) associated with each cluster (D). Top 2,000 cell-type-specific gene markers of the cluster were identified based on a Wilcoxon’s rank sum test comparing with the cluster to SNPs outside of the cluster and enrichment test performed using a hypergeometric test. **D:** UMAP representation of AD SNPs in EBMF from f1 to f6 leiden 1.3. SNPs were colored by clusters and named by the reported gene(s) and displayed its representative SNP or two SNPs. **E-F**: Cell type (F) or selected representative pathways (G) enriched in the top 2,000 genes, the most associated with each SNP cluster. Avg: average, Pval: p-value, FE: Fold.enrichment.

### EBMF-based factors capture distinct SNP influences on AD molecular pathways

Encouraged by the efficient clustering performance using EBMF-based factors, we focused on characterizing each of the EBMF factors for their abilities in capturing the influence of SNPs on diverse AD-associated cellular and molecular cascades. We investigated the first 20 factors, explaining 45% of the total variance (Supplementary Figure 1D). On average, each factor was significantly associated with 15 - 25 SNPs (selection threshold of 1.64 sigma, i.e. 90% confidence interval, to consider as significant SNPs), with most SNPs influencing more than one factor (Figure 3A and Supplementary Figure 2A), suggesting multiple distinct impacts on cell types and pathways. Notably, we observed that the *APOE4* determining variant rs429358 was positively contributing to f4 and f9, while the APOE2 determining variant rs7412 was positively contributing to f2 and f10 (Supplementary Figures 2A). Interestingly, we found that several other variants of the *APOE* locus have independent influence on factors (Supplementary Figures 1E and 2A). Notably, rs405509−T TOMM40|APOE has a leading influence on f5, rs519113−C NECTIN2 has a leading influence on f20, and rs60049679−C APOC1|APOC1P1 has a leading influence on f16 (Supplementary Figures 1E). All these factors are certainly linked to *APOE* locus variants but also to other loci risk variants. For example, the f4 (linked to rs429358 *APOE4* determining variant) being strongly positively influenced by rs13237518−C TMEM106B and rs12590654−G SLC24A4 (Supplementary Figures 2A), suggesting these three variants participating to the same pathological cascade. On the other hand, the variant rs13237518-C TMEM106B, associated with the highest number of DEGs (Supplementary Figure 1C), is also strongly negatively influencing f7 and f10 but positively influencing f1 and f18 (Supplementary Figure 2A), indicative of its putative impact in several independent AD processes.

**Figure 3.**
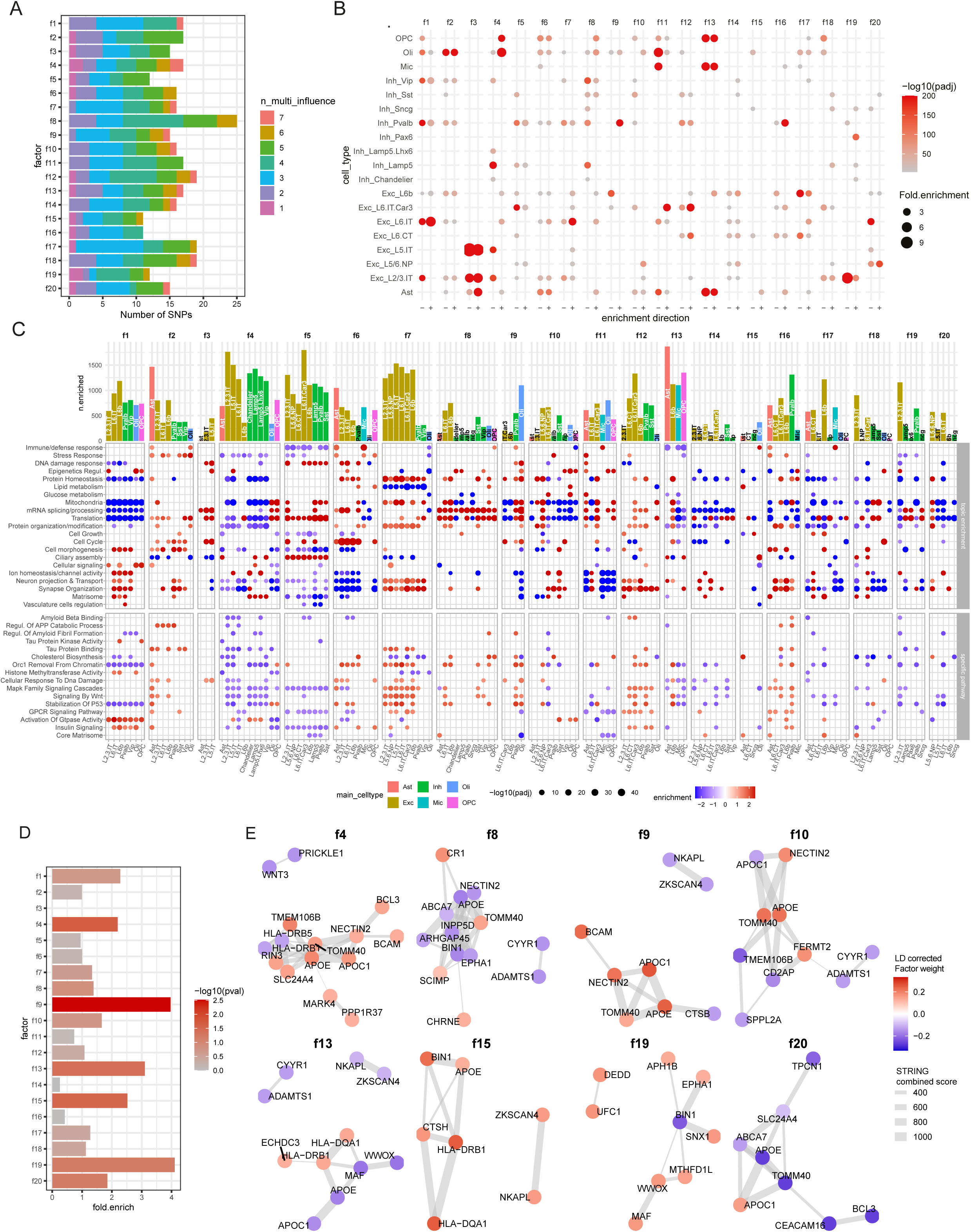
EBMF-based factorization captures functionally coherent SNPs grouped with effects on specific cell types and AD-related molecular pathways. **A**: Number of SNPs associated with each factor. n_multi_influence represents the number of SNPs influencing more than one factor (n_multi_influence = 1 means the SNPs are associated only with one factor). **B:** Cell-type enriched in CtG positively (+) or negatively (-) influenced by each factor, analyzed using a hypergeometric test. **C:** biological topic or selected pathways positively or negatively enriched in CtG loadings for each factor and enriched cell types. **top:** Biological topics positively or negatively enriched, summarized by main category, full details of topics in Supplementary Figure 2B. Enrichment performed using hypergeometric tests on positively or negatively enriched pathways (FDR < 0.01). For the topics, the ‘enrichment’ represents the fold enrichment of the topic category, multiplied by the sign of the direction of the enrichment; for the pathways, the ‘enrichment’ represents the normalized enrichment score (NES) as calculated by fGSEA. **D:** PPI interaction enrichment for the genes of the factor-associated SNPs. The x-axis represents the fold enrichment of the number of interactions in the STRING PPI with combined score > 700 relative to the average number of interactions after degree preserving permutation. **E**: Visualization of top 8 factors (fold enrichment order from D: f19 > f9 > f13 > f15 > f4 > f20 > f10 > f8) with the highest PPI enrichment plotting the reported gene interactions with combined score > 250 for each factor. PPI: Protein protein interaction. Padj: FDR (Benjamini-Hochberg multi-test correction).

We then functionally characterized each of these factors by inspecting the cell types and genes that were most influenced by factors based on their CtG loadings value. Testing enrichment for cell types on the significantly influenced CtG using ORA, we observed that each factor was preferentially associated with distinct cell types (Figure 3B). Performing fGSEA enrichment on the CtG loadings within each cell type, we found that the factor was either positively or negatively influencing a large number of pathways (FDR < 0.01), with several pathways co-influenced by the factors (Figure 3C, Supplementary Figures 2B and 2C, Supplementary Table 6). To summarize this analysis we performed a semantic analysis, grouping each enriched pathway per biological topic and testing for over-representation of these topics in pathways positively or negatively associated with each factor (top panel of Figure 3C and Supplementary Figure 2B). We also displayed enrichment of selected pathways related to AD (bottom panel of Figure 3C). We found that a large majority of these affected processes were reported pathways associated with AD, including immune response, DNA damage response, protein homeostasis/autophagy, mitochondrial activity, synaptic regulation, mRNA splicing, or matrisome signaling^32–34^ (Figures 3B and 3C, Supplementary Figures 2B and 2C). As witness of the coordinated influences captured by these factors, we observed that f3 and f5 were specifically influencing neurons, notably on DNA damage response, cell cycle, translation, and ciliary assembly. F7, which captures notably rs13237518−C TMEM106B and rs147929-A ABCA7 effects (Supplementary Figure 2A), is mainly neuron-specific and positively influences protein homeostasis (especially autophagy), neuron projection, synaptic regulation, MAPK signaling, and tau protein binding, while negatively influences lipid metabolism and cholesterol synthesis pathways (Figure 3C). F13 mostly captures glial specific impacts, being enriched in astrocytes, microglia, and OPCs, with reduction of immune response in these cells (Figures 3B and 3C).

Certain factors have a broader influence on multiple cell types. F4, which captures notably the *APOE4* effect (Supplementary Figure 2A), is enriched in astrocytes, oligodendrocytes, and neurons (Figures 3B and 3C). F4 negatively influences stress response, tau protein kinase activity, and cholesterol biosynthesis in astrocytes, while reducing protein homeostasis (both autophagy and protein ubiquitination), mitochondrial activity, translation, projection and transport in neurons (Figures 3B and 3C). However, f4 increases mitochondrial activity and mRNA processing in oligodendrocytes and OPCs (Figures 3B and 3C, Supplementary Figures 2B and 2C). F2, with leading influence from *APOE2* (Supplementary Figure 2A), is also capturing astrocytes, neurons, and oligodendrocyte impact. F2 increases stress response and protein ubiquitination, MAPK signaling activation, and apoptotic signaling in astrocytes, while increasing APP catabolic process in neurons (Figure 3C and Supplementary Figure 2B). In oligodendrocytes, f2 increases specifically inflammatory, stress, and apoptotic signaling (Figure 3C and Supplementary Figure 2B). These results illustrate how each factor captures different AD relevant molecular cascades or regulation within or across different cell types.

To evaluate how each factor captures already known biological connections between AD risk genes, we tested the interaction enrichment of the reported/candidate causal genes of each factor-associated variants in a protein-protein interaction (PPI) network of all AD risk genes (Figures 3D and 3E). Although the reported genes are not necessarily the causal one, and the PPIs do not integrate all possible direct or indirect gene-to-gene interactions, we identified four factors (f4, f9, f13, and f15) with significant interaction enrichment (p-value < 0.05), and two others factors with suggestive signal (f19 and f20, p-value < 0.1) (Figure 3D), supporting that these factorized risk loci directly cooperate downstream in certain biological process. Among the most enriched, we found that f9, implicating *APOE4*, interact with several other SNPs in *APOE* locus including *TOMM40*, *NECTIN2*, and *APOC1*, and also elsewhere in the genome, in particular, *CTSB*, *NKAPL* and *ZKSCAN4* (Figure 3E and Supplementary Figure 2A). F9 is specifically influencing oligodendrocytes and Inh PvalB, reducing matrisome, lipid metabolism, and synapse organization in oligodendrocytes, while increasing immune response/interferon signaling, and amyloid formation in Inh PvalB (Figure 3C and Supplementary Figure 2B). We also found that f15 (*BIN1*, *HLA-DRB1, HLA-DQA1*, *CTSH, ZKSCAN4,* and *NKAPL*) is positively influencing immune response and ciliary assembly in oligodendrocytes as well as cholesterol synthesis and core matrisome signaling in astrocytes (Figures 3C and 3E).

### Factor-based PGS predicts different neuropathological declines

We questioned whether the factor-based AD risk SNP discrimination could help distinguish the contribution of the SNPs to different AD endophenotypes including amyloid plaque, Tau tangles, as well as most recent findings related to the presymptomatic vulnerable neurons loss^14,16^, epigenome erosion^29^, and increased neuronal genome instability^28^. To test this hypothesis, we computed factor-based polygenic scores (fPGS) using the sum of the factor weights for SNPs multiplied by AD risk allele dose for the SNPs in individuals using an LD whitening method (Figure 1 (4), see Methods). This approach allows us to generate one fPGS for each of the 20 first factors and to measure by individuals for their association with neuropathological outcomes. Using 1,116 individuals from ROSMAP, we first assessed their associations with AD diagnosis and associated clinical outcomes and found significant associations in nine of the fPGSs (f2, f4, f7, f9, f10, f13, f14, f15, and f19) for at least one neuropathological outcome among MCI, AD, disease progression (dcfdx_lv_no6), tau tangles levels (Braak score), Amyloid plaque level (CERAD score), and cognition level (using mini mental state exam, MMSE) (FDR < 0.1; Figure 4A, Supplementary Table 7). Because most of the fPGSs are strongly associated with *APOE4* or *APOE2* genotypes (Figure 4A), we adjusted for the *APOE* genotype to assess which of these clinical associations do not rely exclusively on the *APOE* genotype of the individuals. We found that five out of nine factors are still significant after adjusting for the *APOE* genotype; f7, was strongly and exclusively associated with Braak score, f15, was associated with both Braak score and the semi-quantitative amyloid plaque measurement CERAD score, f4 and f14, were associated with AD diagnosis, and f19, which was negatively associated with MCI (Figure 4B, Supplementary Table 7). We also performed this analysis within *APOE3* homozygotes (Supplementary Figure 3A, Supplementary Table 7), confirming association of f7 and f15 with Braak score (FDR < 0.05), and f4 with disease progression (FDR < 0.1). Interestingly, we noticed that AD-associated f4 and Braak score-associated f7 were both as leading contributors of rs13237518−C TMEM106B but in an opposite manner (Supplementary Figure 2A); the allele contributes positively to f4, whereas negatively to f7, suggesting that a same variant can have both a positive and a negative influence on distinct pathways driving AD pathologies.

**Figure 4.**
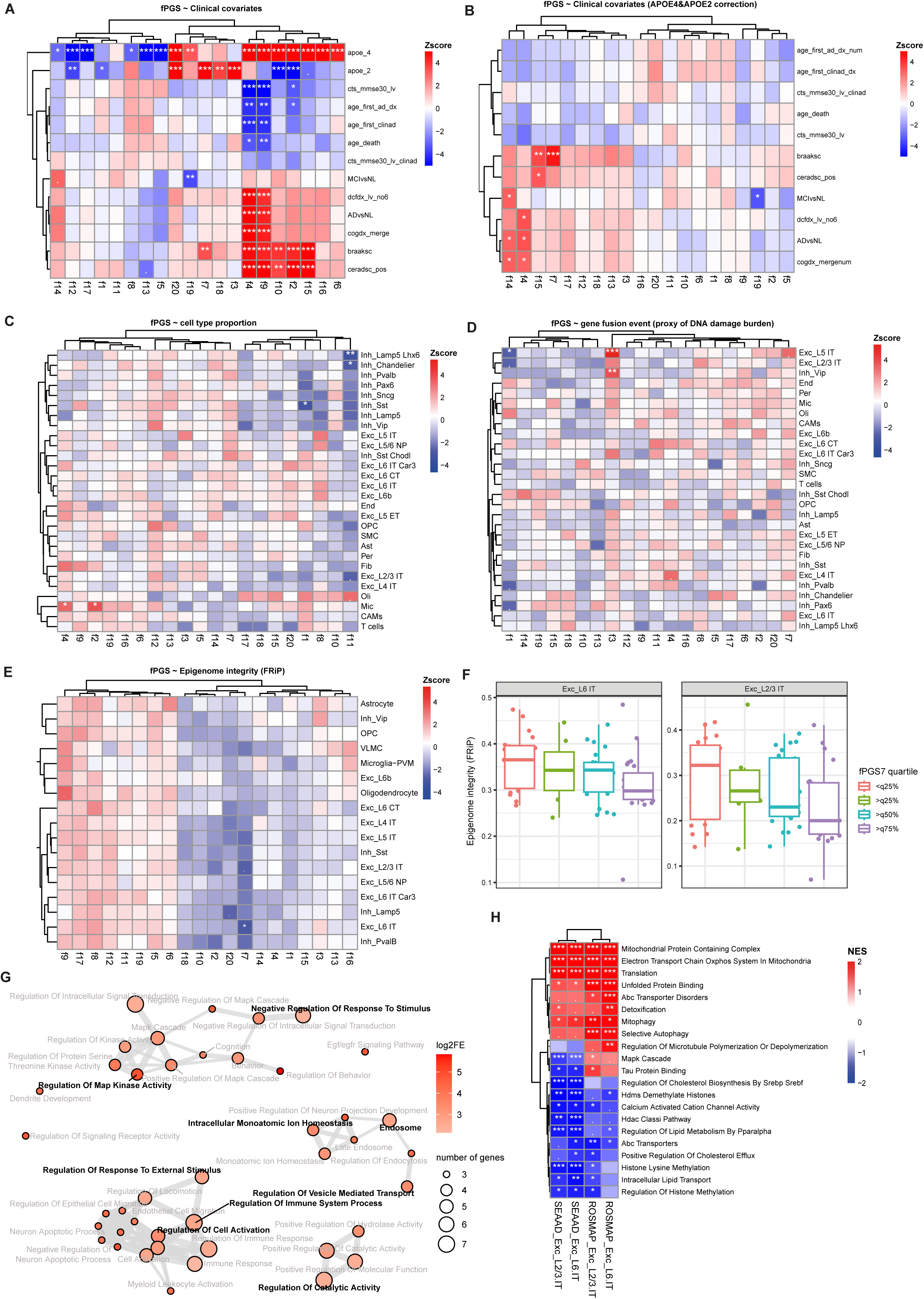
Factor-based PGSs (fPGSs) are associated with specific molecular and neuropathological decline in AD. A-B: fPGSs association with clinical covariates without adjusting for *APOE4* and *APOE2* genotype dose (A), or after adjusting (B). MCI: Mild cognitive impairment; NL: No cognitive impairment; dcfdx_lv_no6: Dementia diagnosis according to ROSMAP clinical codebook, from code 1 (no cognitive impairment) to code 5 (AD and other condition contribution to cognitive impairment), excluding code 6 (others dementia); cogdx_mergenum: consensus cognitive diagnosis, grouping in 3 levels, the cogdx variable from the ROSMAP clinical code: no cognitive impairment (code 1), mild cognitive impairment (code 2 and 3), and AD (code 4 and 5); braaksc: Braak score; ceradsc_pos: CERAD score; cts_mmse30: MMSE test; cts_mmse30_clinad: MMSE test for people with AD; age_first_ad_dx: age-at-first AD diagnosis. Linear regression or logistic regression (ADvsNL and MCIvsNL) adjusting for sex, education, and, if applicable, age-at-death (for the dementia diagnosis, Braak score, CERAD score) or age at last visit (for the MMSE test). **C:** fPGSs association with cell-type proportions from combination of Green *et al.* and Mathys *et al.* ROSMAP DLPFC snRNA-seq datasets. Individuals who were present in the two datasets have been deduplicated by conserving only the individual’s proportions for the dataset with the largest number of cells for the individual. A linear mixed model adjusting for PMI, Sex, age at death, education, number of cells, average cell-type prediction score, with random effect on the datasets. **D:** fPGSs association with gene fusion events (proxy of DNA double-strand break damage burden) detected in snRNA-seq of the Mathys *et al* dataset. A negative binomial regression model adjusting for PMI, sex, education, age-at-death, number of cells, with an offset on sequencing depth. **E:** fPGSs association with fraction of reads in Peaks (FRiP, used as a proxy of epigenome integrity) detected in ROSMAP Xiong *et al.* snATAC-seq dataset. A linear regression model adjusting for PMI, sex, age-at-death, education, and sequencing depth. **F:** Visualization of fPGS7 association with epigenome integrity in excitatory neurons L2/3 IT and L6 IT. Epigenome integrity score is defined as the fraction of reads in peaks (FRiP). **G:** Functional enrichment in reported genes of SNPs significantly contributing to f7 (hypergeometric test with MSigDB GO and CP genesets). **H:** Selected pathways enriched in upregulated or downregulated genes comparing individuals with high (>q75%) vs. low (<q25%) f7 fPGS in ROSMAP and SEA-AD cohorts of Exc L2/3 IT and L6 IT. ***, FDR < 0.001; **, FDR < 0.01; *, FDR < 0.05; ., FDR < 0.1.

To assess if the factors could capture neuronal loss or gliosis, we assessed association of factors with cell-type proportions by combining two ROSMAP DLPFC snRNA-seq datasets^14,26^ including a total of 555 non-overlapping individuals. Because 234 individuals were present in both datasets, we first removed duplicates, considering only the sample for the dataset with the larger number of cells and performed a linear mixed model accounting for the studies’ random effects (Figure 4C). We observed that four factors (f1, f2, f4, and f11) were significantly associated with the cell-type proportion changes (FDR < 0.05); f1 is associated with decrease of the vulnerable somatostatin+ (Sst) inhibitory neurons, which is reported being the first neuronal subtype that is degenerated in DLPFC^16^; f11 is associated with overall decrease of inhibitory neurons, with Lamp5 Lhx6 and Chandelier being statistically significant; and f2 and f4 are associated with increase of microglia (Figure 4C, Supplementary Table 7), indicating factors capture neuronal loss and gliosis. We also performed these associations by merging cells of individuals present in the two datasets instead of excluding one of duplicated samples, and confirmed the associations found in f1, f11, and f4 (Supplementary Figure 3B, Supplementary Table 7). We also observed two other factors (f3 and f8) associated with cell-type proportions; f8 was associated with decrease of inhibitory neurons Lamp5, and f3 was associated with increase of excitatory neurons L5 IT (Supplementary Figure 3B). Interestingly, a pattern of inhibitory neuron loss is observed for f1, f11, and f8, while these three factors are all specifically affecting calcium signaling regulation (Supplementary Figure 2B and Supplementary Figure 3C), concordant with the fact that several reported genes (*SLC24A4*^35^, *SCIMP*^36^, *CHRNE*^37^, *UNC5CL* and *NCK2*-both involved in the calcium-dependant Netrin signaling^38^, *BIN1*^39^, and *ICA1*^40^) of the SNPs associated to these factors are known to be involved in calcium signaling pathways.

DNA double strand break (DSB) damage and following genome instability have been demonstrated as important components of AD neurodegeneration^28^. We examined associations of the fPGSs with DNA damage burden using the number of gene fusion events detected in snRNA-seq as a proxy (Figure 4D, Supplementary Table 7). We observed that f3 was strongly positively associated with gene fusion events in excitatory neurons L5 IT (FDR < 1.10^-8^) and to a lesser extent in inhibitory neurons Vip (FDR < 0.01) (Figure 4D). Consistently, f3 (positively influenced by rs7225151−A ZNF594−DT|SCIMP, rs1800978−G_ABCA1, rs72835061−A CHRNE and rs6489896−C TPCN1), which is specifically associated with transcriptional change in excitatory neurons L5 IT (Figure 3B and Supplementary Figure 2A), affects DNA damage response and repair pathways in addition to chromatin organization, cell cycle, mRNA splicing and catabolism, and ciliary assembly (Figure 3C and Supplementary Figures 3B and 3C). This analysis suggests that specific genetic factors can influence neuronal genome instability and support a link between DSB in excitatory neurons L5 IT and impaired DNA damage response pathways.

Finally, we investigated the association of the fPGSs with epigenome integrity, recently described as associated with cognitive resilience^41^. We used the Xiong *et al.* ROSMAP snATAC-seq dataset^29^ and assessed the fraction of transposition event/reads in open-chromatin region/peaks (FRiP) as a proxy of the epigenome integrity (Figures 4E-4F, Supplementary Table 7). We found that the Braak score-associated f7 (Figure 4B) was associated with decreased FRiP in excitatory neurons L6 IT (FDR < 0.05) and suggestive in L2/3 IT (FDR < 0.1) with a trend in most other neurons (Figure 4E). We confirmed by fPGS quartile analysis in the excitatory neurons (Exc_L6 IT and Exc_L2/3 IT) that the fraction of reads in peaks (FRiP) was inversely correlated as f7 PGS increases (Figure 4F), suggesting the f7 PGS risk well-predicted decreased epigenome integrity. According to the observation, f7 influenced on transcriptomes are mainly enriched in excitatory neurons L6 IT, L2/3 IT, and inhibitory neurons PvalB (Figure 3B). Consistently, f7 affects epigenetic regulation preferentially in Exc L6 IT, but also lipid metabolism, catalytic process/autophagy mitochondrial activity, and MAPK signaling cascade across neurons (Figure 3C, Supplementary Figure 2B and 2C). F7 is influenced by rs13237518−C TMEM106B, rs4147929−A ABCA7, and 14 other SNPs with candidate genes enriched for regulation of MAP Kinase activity and response to stimulus, immune response, ion homeostasis, endosome and catalytic activity (Figure 3G and Supplementary Figure 2A). We assessed if these biological processes and epigenetic changes by f7 PGS can be detected when comparing excitatory neuron gene expressions (Exc_L6 IT and Exc_L2/3 IT) from AD donors with high (> 75% quantile) or low (< 25% quantile) f7 PGS (Figure 4F). We observed indeed in two different cohorts (ROSMAP and SEA-AD) that donors with high f7 PGS had an upregulation in genes of mitochondrial activity and translation, associated with increase of detoxification and selective autophagy activity (especially mitophagy), while a downregulation of cholesterol efflux, lipid metabolism, calcium channel activity and histone modification (Figure 4H, Supplementary Table 7). We also observed dysregulation of MAPK cascade, Tau protein binding, and microtubules polymerization, but with a divergent direction between the two cohorts, which could in part be explained by differences in disease severity (Braak score) (Supplementary Figure 3D). In summary, this analysis suggests a genetically influenced link between increased MAPK signaling, mitochondrial and autophagic activity, reduced lipid metabolism and transport, and reduced epigenome maintenance in excitatory neurons, which could promote tau pathology.

### fPGS-based AD individual clustering distinguishes four clinically relevant genetic risk and resilient subgroups in *APOE* 33 individuals

We finally assessed if the fPGS profiles of AD and MCI individuals are heterogeneous enough to identify clinically relevant genetic risk subgroups. We performed graph-based clustering in the first 20 PC spaces derived from all fPGSs by selecting only AD and MCI ROSMAP donors (N = 768 with 465 AD and 303 MCI). We were able to identify seven donor clusters (cluster 0 - 6) (Figure 5A). Interestingly, these clusters were primarily discriminated by the *APOE* genotype (Figure 5B); cluster 0 was almost exclusively *APOE4* heterozygous or homozygous carriers, cluster 6 was almost exclusively *APOE2* heterozygous or homozygous carriers, and the clusters 1-5 were mostly *APOE3* homozygous carriers, indicating that the fPGSs well-capture the significant and distinct impact of the *APOE* isoforms in AD risk.

**Figure 5.**
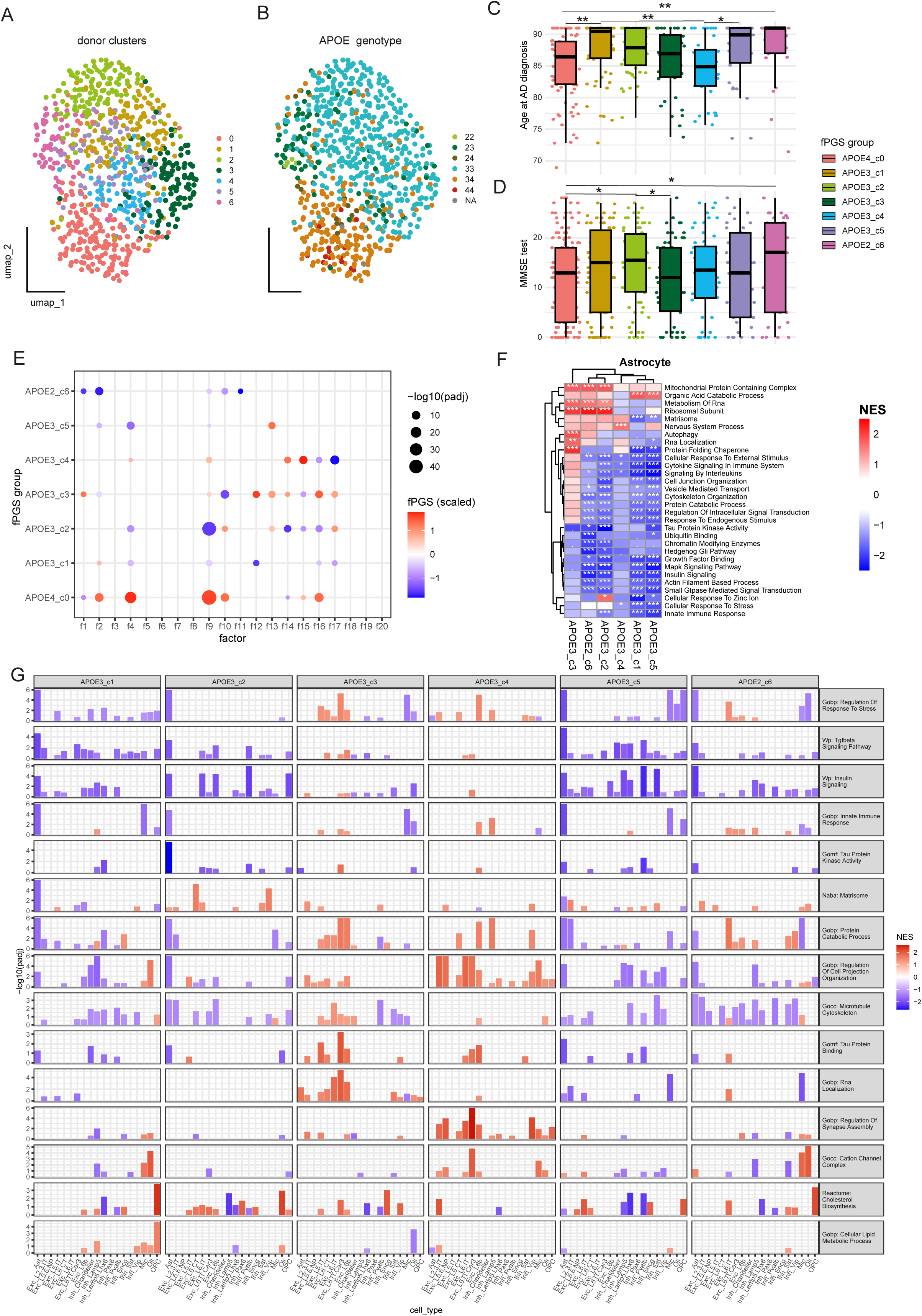
fPGS-based AD donor clustering delineates clinically relevant APOE3 subgroups. A-B: UMAP representation of AD and MCI ROSMAP donor clustering based on their fPGSs (donor cluster 0-6, A) colored by the *APOE* genotype (22, 23, 24, 33, 34, 44, and NA, B). **C:** Association of each AD donor cluster with age-at-AD diagnosis. Linear regression adjusted for sex and APOE4/2 genotype dose. **D:** Association of each AD donor cluster with MMSE. Linear regression adjusted for sex, APOE4/2 genotype dose, and age-at-the MMSE test. **E:** Factors associated with each donor cluster, Wilcoxon test. **F:** Representative pathways enriched in astrocytes when comparing expression in the *APOE3* subgroups or *APOE2* group vs. *APOE4* group (Reference). fGSEA analysis on the DESeq2 stat using MSigDB GO and CP pathways. ***, FDR < 0.001; **, FDR < 0.01; *, FDR < 0.05, ., FDR < 0.25 **G:** Selected enriched pathways in each cell type, showing distinct patterns between the APOE2_c6, resilient APOE3 subgroups (APOE3_c1 and APOE3_c2) and risk (APOE3_c3 and APOE3_c4) groups.

We then investigated how these genetic subgroups were associated with clinical outcomes. We found that these genetic clusters were predicting the age-at-AD diagnosis and cognitive MMSE score of the AD patients (Figure 5C and 5D). As expected, the *APOE4* group (cluster 0, APOE4_c0) is associated with earlier age-at-AD diagnosis as well as a lower MMSE score, while the opposite is observed for the *APOE2* group (cluster 6, APOE2_c6) (Figure 5C). Remarkably, we observed that APOE3_c1 and APOE3_c5 groups were associated with higher age-at-AD diagnosis, with a median age of six years difference for the APOE3_c1 group compared with the APOE3_c4 group, which was lower age-at-AD diagnosis than the APOE4_c0 group (p-value < 0.01, adjusted for sex, education, and *APOE4*) (Figure 5C). On the other hand, we found that the APOE3_c2 group was significantly associated with better cognitive MMSE score, having a 21% greater score compared to the APOE3_c3 group (p-value < 0.05, after adjusting for sex, education, age-at-last visit, and *APOE4*) (Figure 5C). Looking at how each cluster was associated with different fPGSs, we found that f9 was the factor the most differentiating the ‘resilient’ APOE3_c1, APOE3_c2, and APOE3_c5 groups from the ‘risk’ APOE3_c3, APOE3_c4, and APOE4_c0 groups but similar to the ‘protective’ APOE2_c6 group (Figure 5E). Additionally, distinct differences are observed for f4 and f15 being high in the APOE4_c0 and APOE3_c4 group; for f16 being high in the APOE4_c0 and APOE3_c3 group; and for f12 being low in the APOE3_c1 group but high in the APOE3_c3 group (Figure 5E).

At the SNP level, we observed that SNPs that were the most discriminating the cognitively resilient APOE3_c2 group from cognitively-affected risk APOE3_c3 group were rs157580−A APOE (influencing f9, f14), rs1871047−A_NECTIN2 (f9, f10), rs405509−T TOMM40|APOE (f2 and f10), rs584007−G APOC1 (f9, f10), and rs60049679−C APOC1|APOC1P1 (f13, f16) that all reside in *APOE* locus (Supplementary Figure 4A). In contrast, no single SNP was able to discriminate the age-at-AD diagnosis resilient APOE3_c1 and APOE3_c5 groups from the ris k AP OE 3_c4 and APOE 3_c3 groups (S upplementary F igure 4A), suggesting the identification of these resilient groups would not be possible at a single SNP level without the coherent aggregation of genetic signals made possible by the factorization.

Overall, these results indicate that the fPGS-based clustering strategy enables identification of clinically relevant risk and resilient groups in homozygous *APOE3* carriers, predicting disease onset or severity of cognitive decline.

### fPGS-based resilient signals against AD are decreased matrisome and reactivity in astrocytes but increased cholesterol metabolism in OPCs and oligodendrocytes

To further characterize which cell type is involved and what molecular pathways could explain these different clinical outcomes, we performed differential expression analysis and pathway enrichment in each brain cell type comparing *APOE2* or the different *APOE3* subgroups with the *APOE4* group as reference of ‘risk’ molecular profile. We observed a higher number of DEGs (FDR < 0.25 and |log2FC| > 0.25) and pathways (FDR < 0.001) in *APOE2* compared with *APOE4* carrier group (APOE2_c6 vs. APOE4_c0) (Supplementary Figures 4B and 4C, Supplementary Table 8 and 9), consistent with the expected opposite impacts of these two genotypes in AD. *APOE2* has the most DEGs enriched in oligodendrocytes (Oli) followed by excitatory neurons (Exc_L6b) (APOE2_c6 vs. APOE4_c0) (Supplementary Figure 4B). Both *APOE3* resilient group (APOE3_c2 vs. APOE4_c0) and risk group (APOE3_c3 vs. APOE4_c0) display a higher number of DEGs in excitatory neurons (Exc_L6b) (Supplementary Figure 4B). While relatively few DEGs were detected in *APOE3* subgroups, we observed a significant number of pathways enriched by fGSEA across all major brain cell types, indicating a significant number of transcriptional programs diverged among groups and cell types (Supplementary Figures 4B and 4C). We observed that astrocyte was the cell type cumulating the highest number of pathways and functional modules enriched in the resilient APOE2_c6, APOE3_c1, APOE3_c2 and APOE3_c5 groups compared with the *APOE4*_c0 group (Supplementary Figure 4C). Of note, we observed that the risk APOE3_c3 and APOE3_c4 groups had also a large amount of pathways specifically more targeting excitatory neurons (Exc L6b and L6.IT.Car3) but not astrocytes (Supplementary Figure 4C). Comparing the pathways enriched in astrocytes, we observed that all resilient groups (APOE2, APOE3_c1,

APOE3_c2, and APOE3_c5) were specifically associated with reduced gene expressions of response to stress/stimulus, cytokine signaling in immune system, and protein catabolism process, suggesting reduced astrocyte reactivity (Figure 5F). Of note, the resilient APOE3_c1 and APOE3_c5 groups were also specifically associated with reduced matrisome signaling, while the APOE3_c2 group displayed strongly reduced tau protein kinase activity (Figure 5F). It suggests that the AD onset-resilient *APOE3* subgroup has downregulation of matrisome signaling in astrocytes, while the cognitively-resilient *APOE3* subgroup has specific downregulation of Tau protein kinase activity. On the contrary, the risk APOE3_c3 and APOE3_c4 groups showed relatively fewer significantly enriched pathways compared with the resilient groups (Figure 5F). Overall, these results indicate that genetically driven enriched astrocytic matrisome signals and reactivity correlates with worse clinical outcomes.

We further assessed if other molecular signatures can be identified associated with each group across other cell types (Figure 5G, Supplementary Figures 5, Supplementary Table 10). We confirmed that the four resilient groups (APOE2_c6, APOE3_c1, APOE3_c2, and APOE3_c5) were associated with decreased response to stress specifically in astrocytes and were accompanied by decreased TGFbeta and insulin signaling relative to the *APOE4*_c0 group (Figure 5G). These signals were more profoundly reduced in astrocytes but were also found to be reduced in neurons or other glial cells, especially in the resilient APOE3_c1 and APOE3_c5 groups (Figure 5G). The reduced astrocyte reactivity observed in the resilient groups could be explained by their low f4 and f16 fPGS compared with the high fPGS in the APOE4_c0 group (Figure 5E), regarding that both factors influence astrocyte stress or immune responses (Figure 3C). We also found that the strongly reduced tau protein kinase activity in APOE3_c2 group and the strongly decreased matrisome signals in APOE3_c1 and APOE3_c5 were specific to astrocytes (Figure 5G), while the reduction of the protein catalytic process was found both in astrocytes and microglia of the resilient APOE2_c6, APOE3_c2 and APOE3_c5 groups (Figure 5G). On the other hand, the risk APOE3_c3 and APOE3_c4 groups had increased protein catabolic process and cell projection regulation in neurons compared with the APOE4_c0 group, while displaying specific molecular influences; APOE3_c3 excitatory neurons were specifically increased microtubules and tau protein binding process (as well as genes regulating RNA localization), while APOE3_c4 neurons showed increased regulation of synapse assembly and cation channel complex (Figure 5G).

Finally, we observed specifically in the resilient groups a positive enrichment for cholesterol biosynthesis and cellular lipid metabolic process in OPCs and oligodendrocytes (Figure 5G). The increased cholesterol and lipid synthesis pathways were found most predominantly in OPCs of the resilient APOE2, APOE3_c1, and APOE3_c5 groups and in oligodendrocytes of the resilient APOE3_c2 group (Figure 5G). These results are coherent with the observation that the APOE3_c2 group (and to a lesser extent in the APOE3_c1 group) has a very low f9 fPGS compared with the risk groups, especially compared with the APOE4_c0 group (Figure 5E). Indeed, f9 negatively influences cholesterol metabolism in oligodendrocytes (Figure 3C and Supplementary Figures 2B), suggesting that having low f9 fPGS indicates increased cholesterol synthesis. Of note, the increased lipid pathways in the APOE3_c1 group could also be explained by its negative association with f12 fPGS (Figure 5E), regarding that f12 is associated with decreased lipid/cholesterol metabolism in oligodendrocytes (Figure 3C and Supplementary Figures 2B and 2C), again indicating that low f12 PGS predicts increased cholesterol synthesis. Overall, the results of genetically driven resilient signals are associated with decreased matrisome signaling and decreased reactivity in astrocytes but increased cholesterol metabolism in OPCs and oligodendrocytes.

## Discussion

Our study aims at functionally characterizing AD genetic heterogeneity using EBMF by capturing the common influence of a comprehensive panel of AD-associated variants on brain transcriptome. We characterized 20 transcriptional influence patterns (also named ‘factors’) that affects different cellular activities including mitochondrial activity, mRNA processing, synapse regulation, immune and stress response, endolysosomal process, and metabolism of lipids, which are known processes associated with AD^1,4,5,32,33^. These factors are differentially associated with distinct AD neuropathologies, enabling the clinically relevant stratification of AD individuals to genetically predict AD onset and severity.

Using the first six factors, we identified eight AD risk variant clusters with different impacts on brain cell types (Figure 2). One of these clusters specifically influences microglial immune cell activation that responds to lipids and contains SNPs in the *MEF2C* gene locus, the major histocompatibility complex (MHC) class II gene locus, and in *CD33*, *TREM2*, and *ADAM17* loci, each target has an established role in microglial reactivity^42–45^. Another cluster influences synaptic functions, which contain SNPs notably in *SORL1*, *TENM2*, and *FERMT2* loci, also having described roles in synaptic plasticity and APP processing^46–48^. The analysis revealed another SNP cluster containing *CTSH*, *ABCA7*, and *SLC24A4* loci associated with specifically oxidative metabolism and the catabolism of protein aggregates in neurons, and another containing *CR1*, *BIN1*, *ACE*, and *TPCN1* loci, influencing mitochondrial activation, lysosomal pH, and mRNA splicing. These SNP groups are consistent with the known AD pathophysiology and AD patient subtypes previously identified in recent proteomics CSF profile studies, which identified AD subtypes characterized by neuronal hyperplasticity, innate immune activation, and RNA dysregulations^9^.

Generating fPGSs based on each of the first 20 factors, we found 13 factors that were associated with at least one AD endophenotype either at histological, cellular, or molecular level. We observed that f7 fPGS was associated with the Braak score independently of *APOE* genotype (Figure 4B), showing disruption of lipid transport and metabolism, increased mitochondrial activity, selective autophagy, reduced epigenetic integrity, and dysregulated tau activity in excitatory neurons (Figures 3C). This factor implicates risk loci of genes involved specifically in the regulation of cell activation, endocytosis, and MAPK activity, positively associated with rs4147929−A ABCA7, rs112131072−duplication ARHGAP45, rs190982−A MEF2C, rs10498633−G SLC24A4|RIN3 and rs4985556−A IL34 AD risk allele, while negatively associated with rs13237518−C TMEM106B, rs7412−C_APOE (APOE2 defining variant), rs7342692−T PLCG2, and rs5848−T GRN risk allele (Supplementary Figure 2A). Among the implicated genes, *MEF2C* has the most direct role in epigenetics remodeling^49^, by integrating extracellular cues transmitted through MAPK signaling for cellular regeneration, a function that is compromised in aging^50^. Together, this suggests that f7 appears to capture the genetic liability to cell growth signal response and maintain epigenetics integrity needed for neuronal homeostasis. We also found that f3 was associated specifically with increased DNA damage response and cell cycle pathways in L5 IT excitatory neurons (Figure 3C), while the f3 correlates with increased DNA double strand break (DSB)-induced gene fusion and increased proportion of this neuron subtype (Figure 4D, Supplementary Figure 3B). While Dileep *et al.* has found pre-symptomatic elevation of persisting DSB in neurons associated with neurodegeneration^28^, our analysis results further suggest that genomic instability influences in the L5 IT excitatory neuron subtype stronger than other neuronal subtypes and has a genetic predisposition. L5 IT neurons have already been shown to be the most sensitive to DNA damage in mice, which could be linked to their large soma and associated transcriptional demands^51^. This factor is influenced notably by rs7301563−C NTF3, rs1800978-G ABCA1, and rs7155666−G DIO2-AS1|DIO2 (Supplementary Figure 2A). Neurotrophin-3 (NTF3) is playing a critical role to modulate neuronal survival under oxidative stress^52^. ATP-binding cassette transporter A1 (ABCA1) dysfunction leads to impaired cholesterol homeostasis, which amplifies mitochondrial dysfunction and increased oxidative stress^53^. DIO2 converts thyroxine (T4) to active triiodothyronine (T3), which are major regulators of mitochondrial biogenesis, oxidative metabolism, and ROS production^54^, supporting a direct role for these loci in oxidative stress-mediated genomic instability.

Clustering on fPGS profiles surprisingly well-clustered by *APOE* genotypes and identified five distinct *APOE3* homozygous groups in AD patients and MCI individuals (Figures 5A and 5B). Three (APOE3_c1, c2 and c5) of these groups demonstrated enhanced resilience, characterized by either delayed onset of AD or reduced cognitive decline (Figures 5C and 5D), associated with lower matrisome or astrocyte reactivity similar to the resilient APOE2 group (Figure 5F). Previous research has established the significant role of astrocyte reactivity regulation in mediating Tau pathology or resilience^55^. Further, supporting our recent findings nominating astrocyte-driven upregulated matrisome signals in *APOE4* carriers or AD patients^34^, the resilient *APOE3* subgroups showed significant decrease of the matrisome signals (Figures 5F and 5G). Previous reports showed impaired cholesterol biosynthesis detected in oligodendrocytes of *APOE4* carriers and AD patients^56^. Consistently, we found that the resilient groups had elevated cholesterol biosynthesis in OPCs and oligodendrocytes (Figure 5G). Specifically, this pathway was upregulated in OPCs of the AD onset-resilient APOE3_c1 and c5 group and oligodendrocytes of the cognitive-resilient APOE3_c2 group (Figures 5C and 5D). Given the established involvement of oligodendrocytes and OPCs in myelin abnormalities and their recognized contributions to early AD pathogenesis^57^, our findings suggest that upregulated oligodendroglial cholesterol metabolism is a resilient signal against AD.

Previous studies grouped AD risk loci into PGS globally or based on cell-type-specific, co-expression network, or reference-based pathway approaches^13,58–60^. Here, we conducted a transcriptomic data-driven discrimination of all AD risk SNPs to derive clinically relevant genetic groups of AD patients. In contrast to a prior study where global AD PGS predicted AD onset in *APOE4* carriers but not *APOE3* homozygotes^58^, our factor-based PGS (fPGS) approach identified genetic subgroup within *APOE3* homozygotes, exhibiting a median AD onset age 6 years earlier while another subgroup showed a 21% increase in cognitive decline. This highlights that functionally segregating disease risk SNPs substantially improves AD prediction. Compared to a recent study, which built per-cell type PGSs based on gene expression specificity and showed evidence of astrocytic and microglial involvement in AD endophenotype^13^, our findings suggest a contribution from other cell types to AD resilience, specifically identifying oligodendrocytes and OPCs through their upregulation of cholesterol metabolism. These results demonstrate the interest of approaches like EBMF to flexibly capture the combined cellular impact of disease risk loci and provide a more comprehensive picture of cell types and molecular processes involved in the various AD endophenotypes.

To ensure a broad representation of genetic risk, we incorporated a comprehensive panel of AD risk variants identified from multiple GWAS including those with moderate LD. This approach resulted in the inclusion of 24 out of 173 variants located in the *APOE* locus. We found that several variants other than *APOE4* or *APOE2* defining variants were significantly influencing factors (Supplementary Figure 1E) including rs405509−T TOMM40|APOE, (leading variant influencing f2, f10), rs60049679−C APOC1|APOC1P1 (f13, f16), and rs11668327−G TOMM40 (f2, f9, f17) key variants discriminating the resilient APOE33_c2 group from the risk APOE3_c3 and APOE3_c4 groups (Supplementary Figure 4A). R s 405509-G >T at TOMM40|APOE loci has been well-described as an AP OE promoter variant, influencing *APOE* expression with G allele conferring AD protection^61,62^. Rs60049679−C at APOC1|APOC1P1 loci is a variant upstream of *APOC1P1*, which has been associated with C S F Amyloid beta 42 level^63^ and with *APOE* transcript expression in brain^64^. Rs11668327−G>C at the TOMM40 locus is an intronic variant in *TOMM40* and has a known association with blood cholesterol^65^ and Lewy body in AD^66^. These results support the importance of other variants in the *APOE* locus, which together could contribute to *APOE* local haplotype-mediated AD protection.

This factorization by E BMF als o led to deeper fundamental observation. First, our findings showed that a single variant can have both negative and positive influences on distinct AD endophenotypes, as exemplified by rs13237518−C TMEM106B, which negatively influencing epigenome erosion and tau-associated f7, while positively influencing AD progression-associated f4 (Supplementary Figure 2A, Figures 4B and 4E). TME M106B is already known to have a dual role in AD and other neurodegenerative diseases, whereby sufficient expression of TMEM106B is essential for maintaining endolysosomal homeostasis^67,68^, but its excessive expression may lead to lysosomal dysfunction and amyloid fibril formation^69,70^, sugges ting that an allele influencing its expression could have both benefit and drawback depending on the molecular context. More fundamentally, this suggests that a variant can have context-specific effects, for example, being protective for homeostatic processes, while detrimental in challenging conditions . S econd, our findings indicate that two variants, both of which increasing AD risk, can have opposite influences on a particular AD endophenotype, as illustrated by rs13237518−C TMEM106B and rs4147929−A ABCA7, respectively, exerting negative and positive influences on f7 (Supplementary Figure 2A). T his observation supports the evolutionary view that S NP s are naturally selected in a population to mitigate the effect of others and, therefore, keep a biological system robust at a population scale.

### Limitation

As a limitation of this study, we observed only few DEG and SNP influences on microglia despite its known importance in AD genetics and genomics. It may be explained by the lack of power due to their small proportions in the brain, limiting the number of microglia per donor. Due to current technical challenges of vascular cell-type (endothelial and pericytes) detection by single nuclear transcriptome, the study of AD SNP influences on vascular function have not been possible, despite the importance of blood brain barrier breakage in certain subtypes of AD patients and *APOE4* carriers as already shown in previous studies^71^.

Other limitations include the representation of various ancestral populations due to the available gene expression cohort from European ancestry, as well as most of the selected AD variants based on European GWAS studies, limiting the putative representation of our findings in other populations. Furthermore, while we excluded AD risk variants with high LD, more sophisticated finemapping strategies could be applied to refine the list of functionally independent AD risk variants included in the factorization.

### Future Direction

To validate implications of reduced astrocyte matrisome including reactivity and enhanced oligodendroglial lipid metabolism as resilient signals against AD, these findings should be further confirmed in AD resilient populations (individuals with high amyloid plaque and tau tangles burden but low cognitive decline). Further genetic perturbation studies of key regulators of astrocyte matrisome signaling which in part controling reactivity, and/or oligodendrocyte cholesterol production affecting neuronal health and cognitive function using human induced pluripotent stem cell or animal models could be warranted.

## Conclusion

Our study advances our understanding of AD genetics and links with disease heterogeneity, revealing the coordinated impacts of AD risk variants on brain cell-type transcriptome and AD endophenotypes. We report functionally coherent factor-based polygenic scores (fPGSs) predicting specific AD neuropathology and cellular and molecular defects. Notably, one fPGS linking specific AD risk SNPs predicts epigenome erosion and tau pathology, and another fPGS linking another set of AD SNPs points to increased genomic instability in Layer 5 intratelencephalic excitatory neurons. Profiling of these polygenic scores in AD patients have further identified clinically distinct AD subtypes among *APOE3* homozygotes including three resilient subgroups: two subgroups predicting later AD onset, and another subgroup predicting lower cognitive decline. These resilient subgroups are associated with the protective *APOE2* group, with lower matrisome and reactivity in astrocytes, while increasing cholesterol and lipid metabolism in OPCs and/or oligodendrocytes.

## Methods

### Publicly available postmortem brain snRNA-seq data preprocessing

We used the Mathys et al, Religious Orders Study/Memory and Aging Project (ROSMAP) of prefrontal cortex (PFC) single-nucleus RNA sequencing (snRNA-seq) dataset^14^ (syn52293433), containing 372 genotyped individuals with whole genome sequencing (WGS) as training data to perform the differential expression analysis by AD risk variants, the BRAIN Initiative Cell Atlas Network (BICAN)^15^ to define common cell types, and the dorsolateral prefrontal cortex (DLPFC) Seattle Alzheimer’s Disease Brain Cell Atlas (SEA-AD) snRNA-seq data^16^ to annotate the different neuronal subtypes as reference using the label transfer approach from Seurat package^16^.

For SEA-AD snRNA-seq data, gene expression matrix with donors in single-cell level have been downloaded from SEA-AD AWS repository (https://sea-ad-single-cell-profiling.s3.amazonaws.com/ version 2023-07-19) in h5ad single cell format. Donors annotated as ‘Reference’ (young donors lacking cognitive status) have been excluded.

For both ROSMAP and SEA-AD datasets, donor outliers for cell-type distribution have been excluded. Abnormal cell-type distribution was measured by calculating the cell-type proportion deviation from the interquartile range (IQR) and by computing the principal component analysis (PCA) of these deviation matrices. Donors with the first principal component coordinate (eigenvalue) over 3 times away from the IQR were excluded from analysis. For the cell-level quality control (QC), we excluded within each major cell types (excitatory neurons, inhibitory neurons, oligodendrocytes, oligodendrocyte progenitors cells (OPCs), astrocytes, and microglia) if cells with unique molecular identifiers (UMIs) count or number of detected genes over 3 times away from the IQR, and with more than 5% of mitochondrial gene count. To reduce computational burden and keeping only high quality samples for the cell-type label transfer, we downsampled the SEA-AD data selecting the first 12 samples with the lowest average mitochondrial count percentage (average 0.05 % among selected samples).

Before transfer the cell-type labels, the SEA-AD data was normalized using Seurat::NormalizeData(), top 2,000 variable genes were identified using Seurat::FindVariableFeatures(), scaled using Seurat::ScaleData(), and the top 50 PCs was computed using Seurat::RunPCA(). Finally, neuronal subtype labels were transferred using Seurat::FindTransferAnchors() on the 50 computed PCs and Seurat::TransferData().

### AD risk variant selection

To have a broad collection of AD risk loci previously identified across different population, all leading risk variants from the GWAS catalog for traits, ‘Late-onset Alzheimer’s disease’, ‘Alzheimer’s disease,’ and ‘Alzheimer’s disease (late onset)’ have been downloaded (downloaded from https://www.ebi.ac.uk/gwas/ on 2024-05-06, for trait ID EFO_1001870, SI Table 1 and 2). We kept SNPs with p-value < 5e-8 or with suggestive significance < 1e-5 but present at least in two different studies. We also filtered out SNPs in extreme linkage disequilibrium (LD) between each other (R2 > 0.8, keeping only the one with the lowest p-value), and those having less than 5 individuals with the minor allele in the Mathys et al. snRNA-seq dataset^14^, keeping a total of 237 variants for the differential expression analysis. We then annotated the AD risk allele of each of those variants by using Bellenguez et al.^17^ summary statistics from the GWAS catalog as a reference. If SNPs were excluded from the Bellenguez et al. study, the summary statistics were used from Kunkle et al^18^. To annotate each SNP with a neighborhood gene, we used the ‘reported’ gene column of the GWAS catalog summary report. If absent, we used the ‘mapped’ gene column. In case of multiple genes reported/mapped, we displayed only the first two genes.

### Cell-type level differential expression analysis comparing AD risk allele dose

Differential expression analysis in each neuronal or glial cell types was performed using DESeq2 on the pseudobulk matrix (aggregated UMIs count), conserving only individuals with more than 20 nuclei, and genes with at least 1 count per million (cpm) in more than 10% of the individuals. We then tested the risk allele dose effects on gene expressions including only SNPs with at least five individuals in every risk allele dose (0, 1, or 2). To limit high coefficient estimation uncertainty due to low sample size, if a SNPs had less than five homozygotic individuals for the minor allele (*i.e.* the minor allele dose = 2, being represented 4 times or less), we tested only the effect of the *presence* of the minor allele (instead of its dose), *i.e.* we grouped the minor allele homozygotes with the heterozygotes, still requesting at least 5 individuals per group. Differential expressions have been performed, adjusting for the scaled values of post-mortem time interval (PMI), average mitochondrial count percentage, number of cells, age at death, and the first six genotype PCs to account for ancestry difference.

### Empirical Bayes Matrix Factorization (EBMF)-based AD risk variant factorization

We aggregated the z score obtained from differential expression tests of all genes across all cell types and selected variants to generate a variant per cell-type-gene (CtG) matrix.

Before performing the factor analysis, we filtered out SNPs in a 1Mb window to keep SNPs in a low or moderate linkage disequilibrium (LD) (R^2^ < 0.5) or a low or moderate z-score correlation (R^2^ < 0.5). As evidence of independent influence, we also retained SNP pairs if their R^2^ was significantly lower than R^2^ (delta > 0.1). For SNPs that do not reach these criteria, only the SNPs with the lowest p-value in a 1Mb window were kept. This last filtering step allows inclusion of 173 SNPs for the factor-based analysis. We also filtered out the CtG with more than 50% missing z-score (due to more than 50% of the SNPs not tested in this cell type because insufficient number of individuals with enough nuclei).

We performed the factorization by estimating 30 factors (the number of factors K = 30 with K << min(n,p) to ensure a low-rank approximation) using a recently developed Empirical Bayes Matrix Factorization (EBMF) method implemented in the flashier package^19–21^. EBMF flexibly models the data matrix n x p as the product of two low-rank matrices, the factors (F) and the loading (L), of dimension n x K and p x K, with conditions in the modeling based on priors for F and L. Briefly, these priors are estimated based on a user-chosen family distribution, and a variational coordinate-ascent algorithm is used to maximize the variational lower bound (also named as evidence lower bound, or ELBO). Because we expect sparsity into the coordinated effect of the SNPs on the cell activities, we used the ‘point-normal’ mixed distribution for both factors and loadings. Point-normal distributions are sparsity-inducing priors with assumption of normality on the affiliated SNP influences on a specific transcriptional process. Before performing the factorization, we imputed the missing z-score using singular values decomposition (SVD)-based Softimpute package^22^ with rank = 50, lambda = 5, and type = ’svd’ parameter, and then executed an initial singular values decomposition using irlba::irlba() estimating first 30 singular values (parameter nv = 30). EBMF-based factorization was performed using flashier::flash_init() with parameter var_type = 1 and flashier::flash_factors_init() using the singular value decomposition and parameter ebnm_fn = ebnm::ebnm_point_normal. These factor estimations are finally further optimized flashier::flash_backfit() with maxiter = 300.

### EBMF-based SNP clustering performance analysis

We assessed clustering performance and their functional relevance using the EBMF factors comparing with standard PCA approach (PCs obtained from irlba::irlba()) and K-means (using stats::kmeans() with centers = 100 and iter.max = 100 parameter). For EBMF and PCA approaches, The first six EBMF factors or first six PCs have been used to cluster the SNPs using shared nearest neighbor (SNN) graph and leiden modularity optimization^23^. For the K-means center, hierarchical clustering (stats::hclust()) have been used to generate k = 6 and k = 8 clusters using stats::cutree() and match with the number of clusters based on EBMF factors. Silhouette scores have been calculated using cluster::silhouette() for each of these methods. Finally, to assess functional enrichment, CtG changes, which are the most associated with each SNP clusters, have been identified using Wilcoxon rank sum test (stats::wilcox.test()) comparing absolute z-score for the clustered SNPs vs. all other SNPs and the top 2,000 CtGs the most associated based on p-values have been selected filtering out those with nominal p-value > 0.05. The top 2,000 CtGs are then tested for over-representation of specific cell types using hypergeometric test (stats::phyper()), or enrichment of functional pathways using the MSigDB GO and CP gene sets as reference pathways.

### EBMF Factor characterization

Each EBMF factor was first characterized based on the SNPs that were most strongly associated with the factor, considering a SNP significantly contributing to the factor if its absolute value was over 1.64 standard deviation (i.e. 90% confidence interval). We also annotated for what CtGs were associated with each factor by considering significant influences if the CtG had a loading value (weight) for this factor over 1.96 standard deviation (i.e. 95% confidence interval). We characterized these influenced CtGs based on their enrichment for a certain cell type by performing a hypergeometric test. Finally, for each factor and cell types (factor-cell-types) enriched with FDR < 0.001, we performed fgsea::fgsea()^22^ fast-rank gene-set enrichment analysis (fGSEA) using the CtG loading as a gene ranking score against the MSigDB GO and CP gene sets as reference pathways.

### Factor topic analysis

Because all factor-cell-types were enriched for a significant number of pathways (in average 225 per factor-cell-types), we summarized these enriched pathways by topic performing a semantic analysis on the overall enriched pathways with a pretrained language model BERTopic^24^. All enriched pathways in any factor-cell-types at FDR < 0.05 have been included in the semantic clustering analysis. These pathway terms have been projected in the all-MiniLM-L6-v2 embedding space and clustered using bertopic.BERTopic() with language = ‘english’ and hdbscan.HDBSCAN() cluster model with parameter min_cluster_size = 15, and metric = ’euclidean’. It generates 141 clusters of pathways (‘topic’) that were named by the Mistral Pixtral Large Language Model, using the prompt “biological pathways have been grouped by ’topic’ on this csv file, can you find for each of this topic a name which will represent the best the biological theme of these pathways?” in a first round and then improved the answer by prompting “for each of this topic can you find a name more concise, like topic 59 should be ’Autophagy & Organelle Degradation’ instead of ‘Autophagy, Regulation, Mitophagy, Autophagosome, Mitochondrion’”. Then, each factor-cell-type was tested for over-representation for each of these topics performing an hypergeometric test on the pathways positively or negatively associated (FDR < 0.05) to the factor-cell-types.

### PPI enrichment

Each of the SNPs contributing to each factors were tested for their downstream functional relationship by testing how well each of the reported genes for the SNPs (using REPORTED GENE(S) column in the GWAS catalog summary table (SI Table 2), or the MAPPED_GENE gene column if the first is blank) were interconnected in STRING-db Protein Protein Interaction (PPI) network (9606.protein.links.detailed.v12.0), using combined_score threshold of 700. For each factor, we performed a degree-preserving randomization via double-edge swaps (Markov-chain Monte Carlo), testing the number of interconnections (edges) for the SNP-associated AD risk genes present in the PPI database compared with the number of interconnections after shuffling all edges 10 times among all reported genes included in this study, and permuting 2,000 times.

### Factor-based polygenic score (fPGS) construction

For each factor, we constructed a polygenic score (PGS) weighting AD variants based on their weight in the factor and applying an LD-whitening transformation using the inverse square root method to avoid double-counting genetic signals due to LD^25^. Starting with the variant with the highest absolute factor weight, we first estimated the SNP–SNP correlation matrix R for variants within 1 Mb windows calculated using the full WGS ROSMAP cohort (n = 1,116 individuals). We then computed the inverse square root of the LD matrix (R⁻¹ᐟ²) by eigendecomposition and replaced the original factor weight w to LD-corrected factor weights, ŵ:

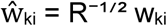

where R denotes the variant-variant pearson correlation matrix, and w_ki_ is the weight of variant i in factor k.

This transformation projects the variant loadings onto an orthogonal basis, effectively removing linear dependencies due to LD, while preserving the total variance explained by the set of variants. The final factor-based PGS (fPGS) were then calculated as:

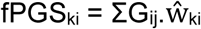

where Gᵢⱼ denotes the AD risk allele dosage of variant i in individual j, and ŵ_ki_ is the transformed corresponding LD-corrected factor weight for variant i.

### fPGS association with clinical covariates

We used the entire WGS ROSMAP cohort (n = 1,116 individuals) to perform this association. Each following AD-associated clinical covariates have been tested: age-at-death, CERAD score, Braak score, clinical cognitive status (excluding ‘others dementia’), Mini-Mental State Examination (MMSE) level, and final cognitive status (examined by a neurologist at death), which have been divided into three levels: non cognitively attained (NL, code 1), mild cognitive impairment (MCI, code 2 and 3), and AD (code 4 and 5). We also tested specifically AD vs. NL and MCI vs. NL. We tested association of those clinical covariates with each of the first 20 fPGS using linear regression (except for the binary variable AD vs. NL and MCI vs. NL, where we ran a logistic regression using stats::glm(,family=binomial)), adjusting for sex, education, and, if applicable, age-at-death (for the clinical diagnosis, Braak score, CERAD score) or age at last visit (for the MMSE test) with following formula:

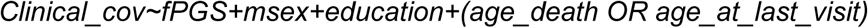

Associated t value/z score for the fPGS was then extracted, and FDR was calculated by correcting for the number of factors. We also tested for *APOE4* and *APOE2* allele dose association in the same way to assess the dependency of each fPGSs to *APOE* genotype.

### fPGS association with cell-type proportions

To assess association with cell-type proportion, we integrate the Mathys et al.^14^ dataset (syn52293433) and the Green et al.^26^ ROSMAP snRNA-seq dataset (syn31512863) to maximize the number of samples and mitigate possible batch effects (n = 555 individuals). We reprocessed these two datasets from the fastq files to have the same QC procedure to reduce the technical variabilities in cell-type proportion estimates. The UMI count matrix have been generated from each fastq file using simpleaf quant^27^ with index generated from the 10XGenomics refdata-gex-GRCh38-2020-A hg38 genome and GTF reference using parameters ‘--chemistry 10xv3’, ‘--resolution cr-like’, ‘--unfiltered-pl’, and ‘--expected-ori fw’. UMI count matrix is loaded in R with fishpond::loadFry(outputFormat = ’snRNA’) to account for both exonic and intronic UMIs per gene, and cells are filtered out if less than 1,000 UMIs or 800 genes detected, or more than 10% of mitochondrial count. For the Green et al. dataset containing pooled samples by libraries, samples are demultiplexed using the ROSMAP_snRNAseq_demultiplexed_ID_mapping.csv (syn34572333). Cell-type labels are then transferred to the SEA-AD neurons annotated by the DLPFC reference generated previously using FindTransferAnchors() and TransferData(), as previously described in section ‘Publicly available postmortem brain snRNA-seq data preprocessing’. Cells with prediction.score.max < 0.6 for cell-type labels were filtered out, and this score was further used as QC metrics for the cell integrity. Because these two datasets have a significant number of duplicated individuals (n = 234 individuals overlapped in the two datasets), we removed one of the duplicates with the lowest number of total cells. Finally, cell-type proportion associations with fPGS have been tested using a mixed model using lme4::lmer() by adjusting for random effects to account for the dataset variabilities and including PMI, sex, age-at-death, education, total number of cells, and average cell-type prediction score as a fixed effect using the following model:

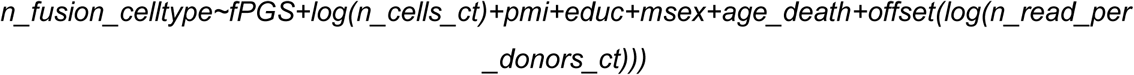

We also performed this analysis using a different approach to consider the duplicated individuals. In this alternative approach, nuclei from the same individual present in the two datasets have been grouped, and cell-type proportions have been calculated afterward, preventing the sample exclusions. A linear regression model has then been used to adjust for PMI, Sex, age-at-death, education, number of cells, and average cell-type prediction score as cell quality score.

### fPGS association with somatic gene fusion event

We downloaded the preprocessed number of gene fusion events per cell (https://personal.broadinstitute.org/cboix/dileep_boix_et_al_data/Data/human-snRNA-seq/) from Dileep et al.^28^ and aggregated the count per individual and cell type. We then used a negative binomial regression model using MASS::glm.nb() with an offset term for the number of reads captured per sample, and correcting for the number of cells, PMI, education, sex, and age-at-death with the following formula:

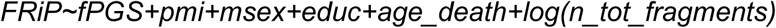

### fPGS association with epigenome integrity

Xiong et al.^29^ ROSMAP snATAC-seq fragment files (syn52335505) have been processed using Signac^30^. Briefly, peaks from the fragments files are first called for each individual using Signac::CallPeaks() with macs3 (v3.0.1)^31^. Peaks identified in each individual are then combined across all individuals using GenomeRanges::reduce(). The peak count matrix for each individual across all nuclei was then generated based on these combined peaks reference using Signac::FeatureMatrix(), and nuclei are conserved if more than 500 counts in peaks. To annotate the cell type, we first transferred cell-type labels from the snRNA-seq SEA-AD DLPFC reference on the samples with the highest quality metrics. The fractions of reads in peaks (FRiP) and TSS Enrichment (using Signac::TSSEnrichment() and EnsDb.Hsapiens.v86 annotations) were calculated, and the top 20% samples were selected based on a composite score combining the fraction of reads in peaks (FRiP; +1, if the individual is in the top 20% samples with the highest FRiP), median number counts in peak per cells (+1 if in top 20% samples with the highest count), and TSS enrichment (+1, if in the top 20% samples with the highest TSS enrichment). These high quality samples were merged, and nuclei were further filtered to conserve only those with number of reads in peaks greater than 1,000 but smaller than 100,000, FRiP > 0.25, nucleosome signal (computed with Signac::NucleosomeSignal()) < 4 and TSS.enrichment > 2, resulting in preserving 64,651 nuclei. The gene activities (i.e. overall number of reads around genes) were then calculated using Signac::GeneActivity(), and snRNA labels were transferred using Seurat::FindTransferAnchors() with reduction = ’cca’ and dims = 1:40, followed by Seurat::TransferData() using for a weight.reduction parameter, a precomputed latent semantic indexing (LSI) reduction (computed using default Signac::RunTFIDF() and Signac::RunSVD()). These cell-type labels were then transferred to all samples using FindTransferAnchors() with parameters reference.reduction = “lsi”, reduction = “lsiproject”, dims = 2:30 and MapQuery() with reference.reduction = “lsi”. The individuals with ethnicity ‘Black or African American’ have been excluded as well as donors with abnormal cellular distribution using the same approach as with the snRNA-seq data. The sample ‘D19-12532’ was excluded because it had a very different chromatin profile for Exc subtypes compared to other samples (suspecting a brain region mismatch), as well as ‘D19-12545’ having outlier profile for oligodendrocytes. In total, 9 out of the total 92 samples were excluded.

Before computing the final FRiP per sample as a proxy of the epigenome integrity, we increased a resolution of peak identifications by calling peaks per cell type and group of donors, with an assumption that some donors could have specific cell states and thus open chromatin region profile may not be detectable (ratio signal/noise being too low). To cluster donors according to their chromatin profile, we first call peaks using Signac::CallPeaks() within each cell type, selecting only the top 20% previously selected and QC-ed samples. We then used the resulting peak count matrix (aggregated per sample) to cluster the samples using a LSI dimension reduction with Seurat::FindNeighbors() on the first six dimensions and Seurat::FindClusters() with algorithm = 3 and resolution = 1.2. We finally performed the final peak calling per sample cluster using Signac::CallPeaks() within each cell type and combined all peaks identified using GenomeRanges::reduce(), resulting that 491,503 unique peaks were identified, while only 367,242 were identified in the Xiong et al. study^29^. We notably identified around 20,000 peaks in astrocytes that were specific to one donor cluster (and similar observations found in Excitatory L5 IT and Excitatory L6b), demonstrating interest to perform peak calling per group donor and cell type rather than globally.

Finally, we computed the FRiP per individual and cell type and used a linear model to test association between FRiP and fPGS, correcting for PMI, sex, age-at-death, education, and total number of fragments with the following formula:

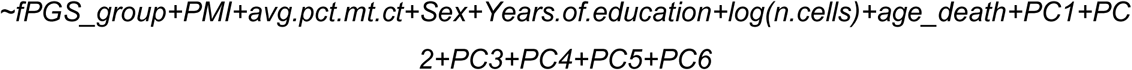

### fPGS-based AD and MCI donor clustering

PCA has been performed on AD and MCI donors for the fPGS generated for the 30 computed factors, and the first 20 PCs have been used to cluster the donors using Shared nearest graph using Seurat::FindNeighbors() and Seurat::FindClusters() with resolution = 1, as well as for generating the UMAP using Seurat::RunUMAP() using n.neighbors = 10.

### Differential expression analysis and pathway analysis of fPRS groups

We performed a cell-type level differential expression analysis comparing fPGS > q75% vs. fPGS < q25% or comparing any two different fPGS-based subgroups using DESeq2 on the pseudobulk matrix in Mathys et al. and SEA-AD DLPFC snRNA-seq data^14,16^, correcting for PMI, mitochondrial percentage, sex, education, number of cells, age-at-death, and the first six genotype PCs using the formula:

*~fPGS_group+PMI+avg.pct.mt.ct+Sex+Years.of.education+869 log(n.cells)+age_death+PC1+PC2+PC3+PC4+PC5+PC6*

We then performed gene-set enrichment analysis using fgsea::fgsea() on the resulting DESeq2 statistics to identify enriched MSigDB GO or CP pathways in up- or down-regulated genes.

## Data availability

The datasets analyzed for the current study are available in Synapse. snRNAseq data are accessible under the accession codes from their original publications syn52293433, syn26223298, syn31512863, for Mathys et al., Gabitto et al. (SEA-AD) and Green et al., respectively, under controlled use conditions due to human privacy regulations.

The code used for this study will be made available upon publication at [github link]

## Abbreviations

AD: Alzheimer’s disease
APOE2/3/4: ε2, ε3, or ε4 allele of Apolipoprotein E
BICAN: BRAIN Initiative Cell Atlas Network
CSF: Cerebrospinal fluid
CT: Corticothalamic
CtG: cell-type-gene (expression of a gene in a cell type)
DLPFC: dorsolateral prefrontal cortex
EBMF: Empirical Bayes Matrix factorization
Exc: Excitatory neurons
IT: intratelencephalic
FDR: False discovery rate
fPGS: Factor-based Polygenics scores
fGSEA: fast-rank Gene-set Enrichment Analysis
FRiP: Fractions of reads in peaks
GWAS: Genome-wide association studies
IQR: Interquartile range
Inh: Inhibitory neurons
IT: intratelencephalic
Exc L2/3/5/6: Excitatory neurons of cortical layer 2/3/5/6
LD: Linkage disequilibrium
LSI: Latent semantic indexing
MCI: Mild cognitive impairment
MMSE: Mini-Mental State Examination
NES: Normalized enrichment score
NL: Non cognitively attained
NP: Near projecting
OPCs: Oligodendrocyte progenitors cells
ORA: Over-representation analysis
PCA: Principal component analysis
PFC: Prefrontal cortex
PGS: Polygenics scores
PMI: Post-mortem time interval
PPI: Protein Protein Interaction
QC: Quality control
ROSMAP: Religious Order Study and Memory and Aging Project
SEA: Seattle Alzheimer’s Disease Brain Cell Atlas (SEA-AD)
SNN: Shared nearest neighbor
SNPs: Single nucleotide polymorphisms
snATAC: Single-nucleus Assay for Transposase-Accessible Chromatin using sequencing
snRNA-seq: Single-nucleus RNA sequencing
SVD: Singular values decomposition
TSS: Transcription start site
UMIs: Unique molecular identifiers
WGS: Whole genome sequencing

## Acknowledgements

The results presented here are based on data obtained from the AD Knowledge Portal (https://adknowledgeportal.org). These data would not be possible without the participation of research volunteers and the contribution of data by collaborating researchers. ROSMAP data were provided by the Rush Alzheimer’s Disease Center, Rush University Medical Center, Chicago. Whole genome sequencing was supported through funding by NIA grants U01AG61356 (whole genome sequencing); Mathys et al. snRNA-seq data and the snATAC-seq data collection was supported through the Cure Alzheimer’s Fund, NIH grants AG058002, AG062377, NS110453, NS115064, AG062335, AG074003, NS127187, MH119509, HG008155 (M.K.), RF1AG062377, RF1 AG054321, RO1 AG054012 (L.-H.T.) and the NIH training grant GM087237 (to C.A.B.). Green et al. snRNA-seq data collection was supported through NIH grants U01AG061356 (De Jager/Bennett), RF1AG057473 (De Jager/Bennett), and U01AG046152 (De Jager/Bennett) as part of the AMP-AD consortium, as well as NIH grants R01AG066831 (Menon) and U01AG072572 (De Jager/St George-Hyslop). SEA-AD study data were generated from postmortem brain tissue obtained from the University of Washington BioRepository and Integrated Neuropathology (BRaIN) laboratory and Precision Neuropathology Core, which is supported by the NIH grants for the UW Alzheimer’s Disease Research Center (P50AG005136 and P30AG066509) and the Adult Changes in Thought Study (U01AG006781 and U19AG066567), and by NIA grant U19AG060909.

We thank Gao Wang, Zining Qi, and the FunGen AD xQTL working group for the collaborative training and important discussions on robust statistical methods for omics data modeling, which have inspired this study. We also thank Jason Willwerscheid, who developed the Empirical Bayes Matrix Factorization (EBMF) framework on the flashier package, for his insight on the EBMF methods and applications. We would also thank Alec Candib for his assistance with the initial QC and cleanup of ROSMAP data during the lab rotation, and Yun Shen for the research computing at Boston University Information Services & Technology.

## Ethics declarations Competing interests

JTCW serves on the scientific advisory board of NeuCyte, Inc, and has consulted for CareCureSystems. The authors declare no competing interests.

## Ethics approval and consent to participate

Detailed information on ethical approvals and participant consent for the ROSMAP snRNA-seq data^14,26^, whole-genome sequencing data^72^, and SEA-AD data^16^ has been reported previously. The ROSMAP studies were approved by the Institutional Review Board (IRB) of Rush University Medical Center, and the whole-genome sequencing studies were approved by the IRB of Massachusetts General Hospital under protocols 2015P000111 and 2019P001879.

## Figures legends

**Supplementary Figure 1. AD risk variant selection and DEG analysis. A**: Selected AD risk variants by various GWAS studies. The x-axis represents the AD GWAS SNPs (format rsid-risk_allele reported_cene(s)) selected across 23 GWAS studies if (i) genome-wide significant (p < 5e-10) or suggestive (p < 1e-5) in at least two studies, (ii) having minor allele count > 5 in the snRNA-seq dataset, and (iii) R^2^ < 0.5 with any other SNPs or having independent influence on transcriptome (see Method). The y-axis represents -log10(p-value) for their associations with AD in the study (truncated to 100). **B**: Distribution of R^2^ and R^2^ for each GWAS SNPs with the most highly correlated SNPs in the 1Mb window for SNPs passing QC (TRUE) and the rest (FALSE). **C**: Number of DEGs (FDR < 0.05 and |log2FC| > 0.25) detected per cell type for each variant, for the top 50 variants with the highest number of DEGs. **D**: Proportions of variance explained for each factor of EBMF on the SNP-gene-cell type differential expression matrix. **E**: Factor weights for the AD risk variants in the *APOE* locus (+/-1Mb of *APOE*). *, weight above 1.64 sigma.

**Supplementary Figure 2. EBMF-based factor characterization. A**: SNPs associated with each factor. The SNP is considered as “associated to a factor,” if its absolute weight is above the 1.64 standard deviation. Dotsize represents absolute weight, while color represents the positive (red) or negative (blue) weight after correcting for LD with the rest of SNPs. *APOE4* and *APOE2* determining SNPs are in red and blue, respectively. **B**: Biological topic over-represented in positively or negatively enriched pathways in CtG loadings for each factor within each factor associated cell types. Signed log2FE: log2 of the fold enrichment (FE) for topic over-representation in enriched pathways, represented in blue if the topic enrichment is within negatively enriched pathways (NES < 0) or in red if positively enriched pathways (NES > 0). **C**: MSigDB representative pathways enriched across all factors by each enriched cell type, enrichment analysis performed on the CtG loading value in each factor using fGSEA.

**Supplementary Figure 3. fPGS associations with specific molecular and neuropathological decline in AD. A**: fPGS associations with clinical covariates within *APOE3* homozygotes (n=675), using the same approach as Figure 3A. **B**: fPGSs association with cell-type proportion fully merging Green *et al.* and Mathys *et al.* ROSMAP snRNA-seq datasets^14,26^. Nuclei from the same individual present in these two datasets have been merged, accounting for one cell-type proportion. A linear regression model adjusting for PMI, Sex, age at death, education, number of cells, and average cell type prediction score. **C**: Factors enriched for topic ‘Calcium Ion Transport & Signaling Regulation’. ***, FDR < 0.001; **, FDR < 0.01; *, FDR < 0.05; ., FDR < 0.1. **D**: Braak score and CERAD score distribution in SEA-AD and ROSMAP AD donors. ****,: p<0.0001, Wilcoxon rank sum test. ns, not significant.

**Supplementary Figure 4. fPGS-based AD patient clustering characterization. A**: SNPs associated with each cluster. Wilcoxon rank sum test on the cluster donors’ SNP dose profile compared with donors of other clusters, FDR < 0.01 and |log2FC| > 0.25). **B**: Number of DEGs (FDR < 0.25 and |log2FC| > 0.25) comparing the different donor clusters (c1-c6) with the *APOE4* cluster (c0) in each cell-type. **C**: For each cell-type, number of pathways (FDR < 0.001) and functional groups enriched in different groups compared with the APOE4 group (reference) using fGSEA. Functional pathway groups are defined by grouping them based on the jaccard similarity graph with edges between pathways if Jaccard similarity > 0.2 and using leiden algorithm with resolution = 1. D-E: Representative selected pathways enriched among the group in excitatory neurons L6b (D), or in inhibitory neurons Sst (E)

**Supplementary Figure 5. Functional enrichment in the different AD patient clusters compared to the *APOE4* cluster. A-C**: Representative selected pathways enriched among the donor groups (referenced to the *APOE4* group) in excitatory neurons L6b (A), inhibitory neurons Sst (B), oligodendrocyte (C), OPC (D), microglia (E).

## Tables Legends

**Supplementary Table 1.** GWAS studies and number of variants included in this analysis. ‘n_SNPs’ is the number of leading variants identified in this study, which have been tested for differential expression analysis, and ‘n_SNPs_passQC’ is the number of these variants that pass QC to be included in the factorization analysis.

**Supplementary Table 2.** AD risk variants selected in this study. ‘pass_QC’ if the variant pass the LD and z-score R2 QC to be included in the factorization

**Supplementary Table 3.** Differentially expressed genes (FDR < 0.05 & |log2FC| > 0.25) per variant and cell type.

**Supplementary Table 4.** Variant clustering based on the six first EBMF factors.

**Supplementary Table 5.** Enriched pathways (FDR < 0.25) in the top 2,000 CtG gene influence markers of each SNPs cluster.

**Supplementary Table 6.** Pathways enriched (FDR < 0.05, fGSEA) either positively or negatively for each factor CtG loadings, and topic enrichment in these pathways.

**Supplementary Table 7.** Null hypothesis testing for fPGS association with clinical, cellular or molecular outcomes.

**Supplementary Table 8.** Pathways enriched (FDR < 0.05, fGSEA) in differential expression results comparing f7 PGS > q75% vs. < q25% donors Exc_L2/3 IT and Exc_L6 IT transcriptome in ROSMAP and SEA-AD cohorts.

**Supplementary Table 9.** Differentially expressed genes (FDR < 0.25 & |log2FC| > 0.25) of the fPGS-based donor clusters compared to the APOE4_c0 cluster in each cell type.

**Supplementary Table 10.** Pathways enriched (FDR < 0.05, fGSEA) in fPGS-based donor clusters compared to the APOE4_c0 cluster in each cell type.

