## Supplementary Figures for "Factorization of Alzheimer’s disease genetic risk influences allow patient stratification, predicting disease onset, cognitive decline, and cell-type specific responses"

### Supplementary Figure 1

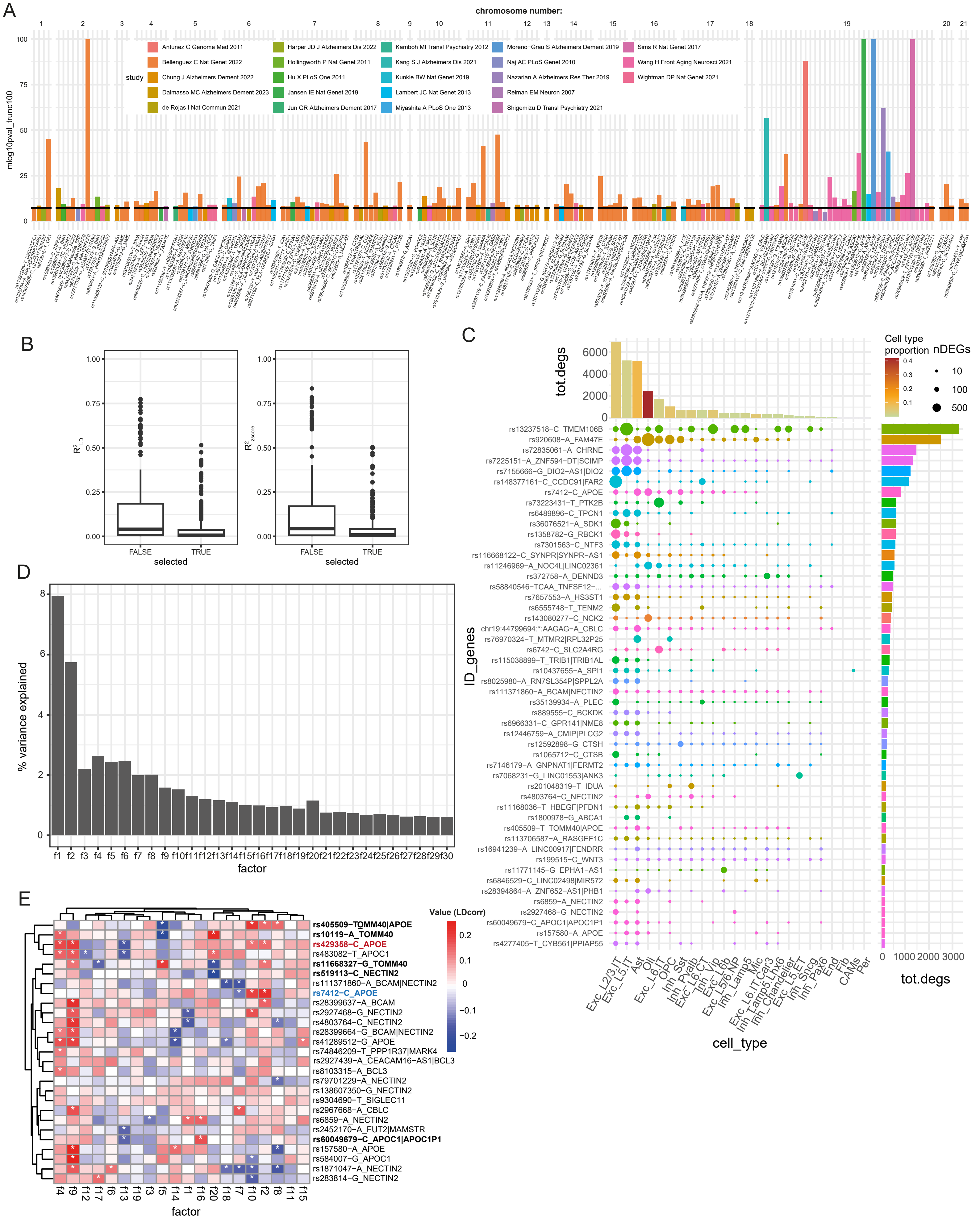

Supplementary Figure 2

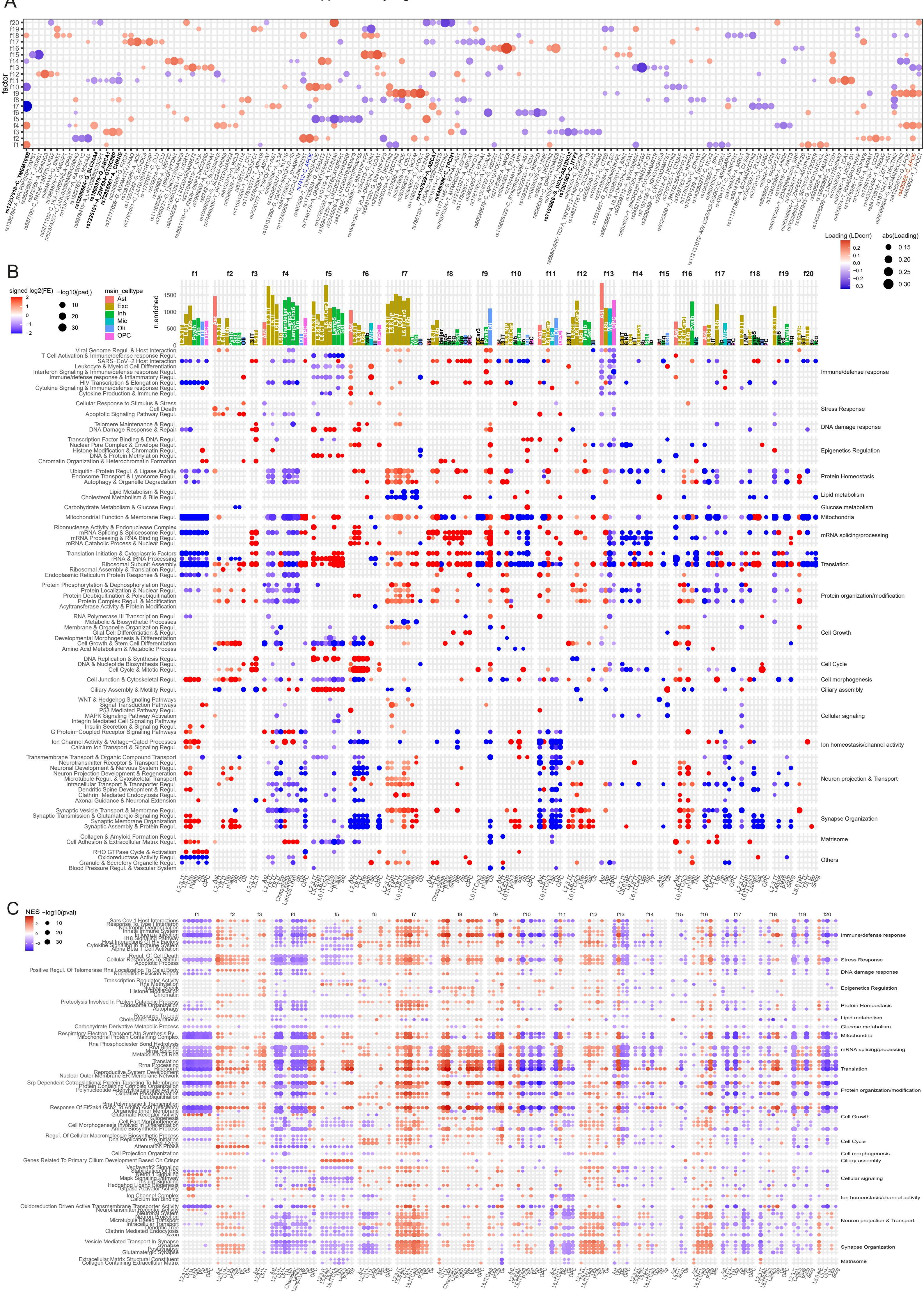

### Supplementary Figure 3

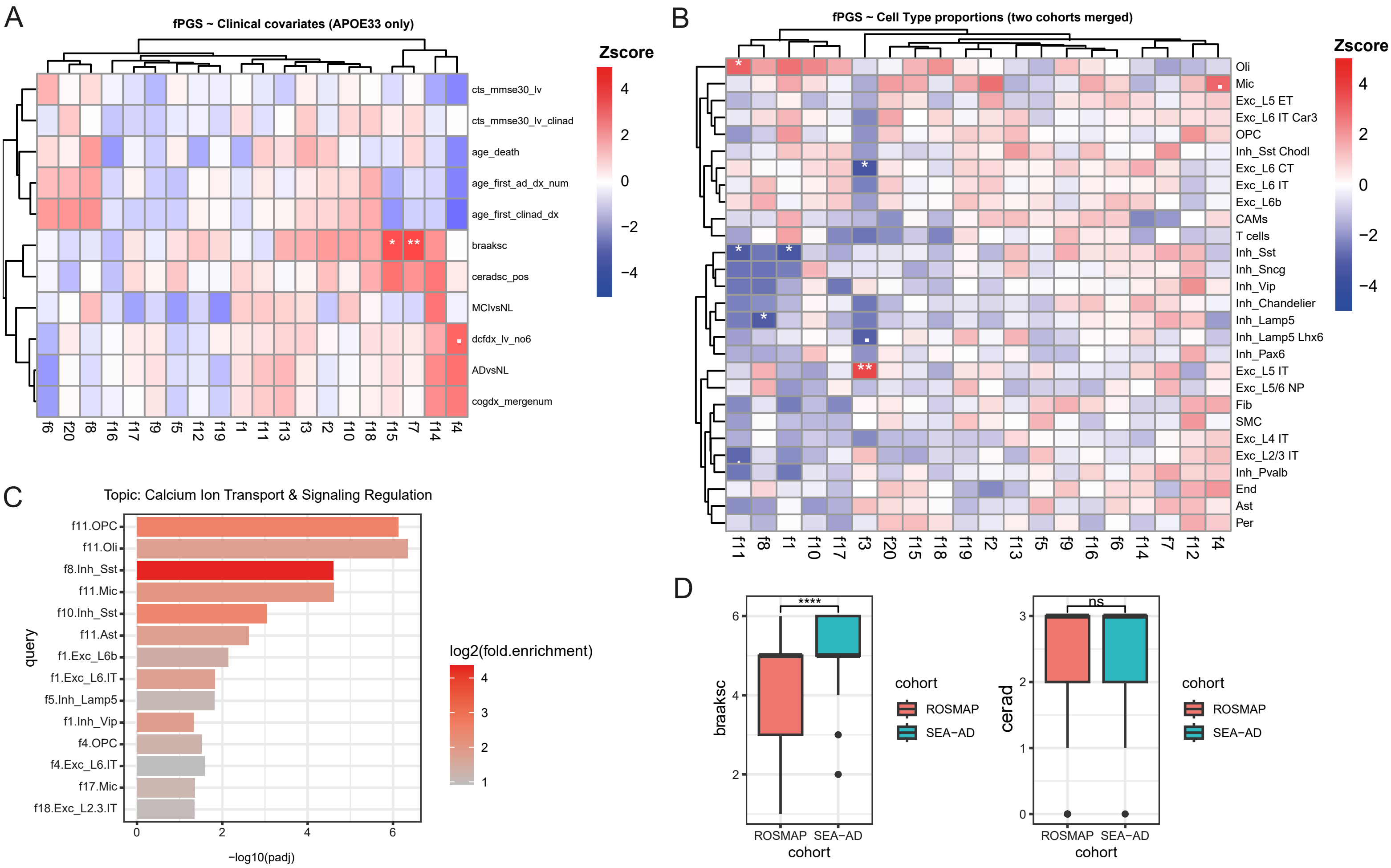

Supplementary Figure 4

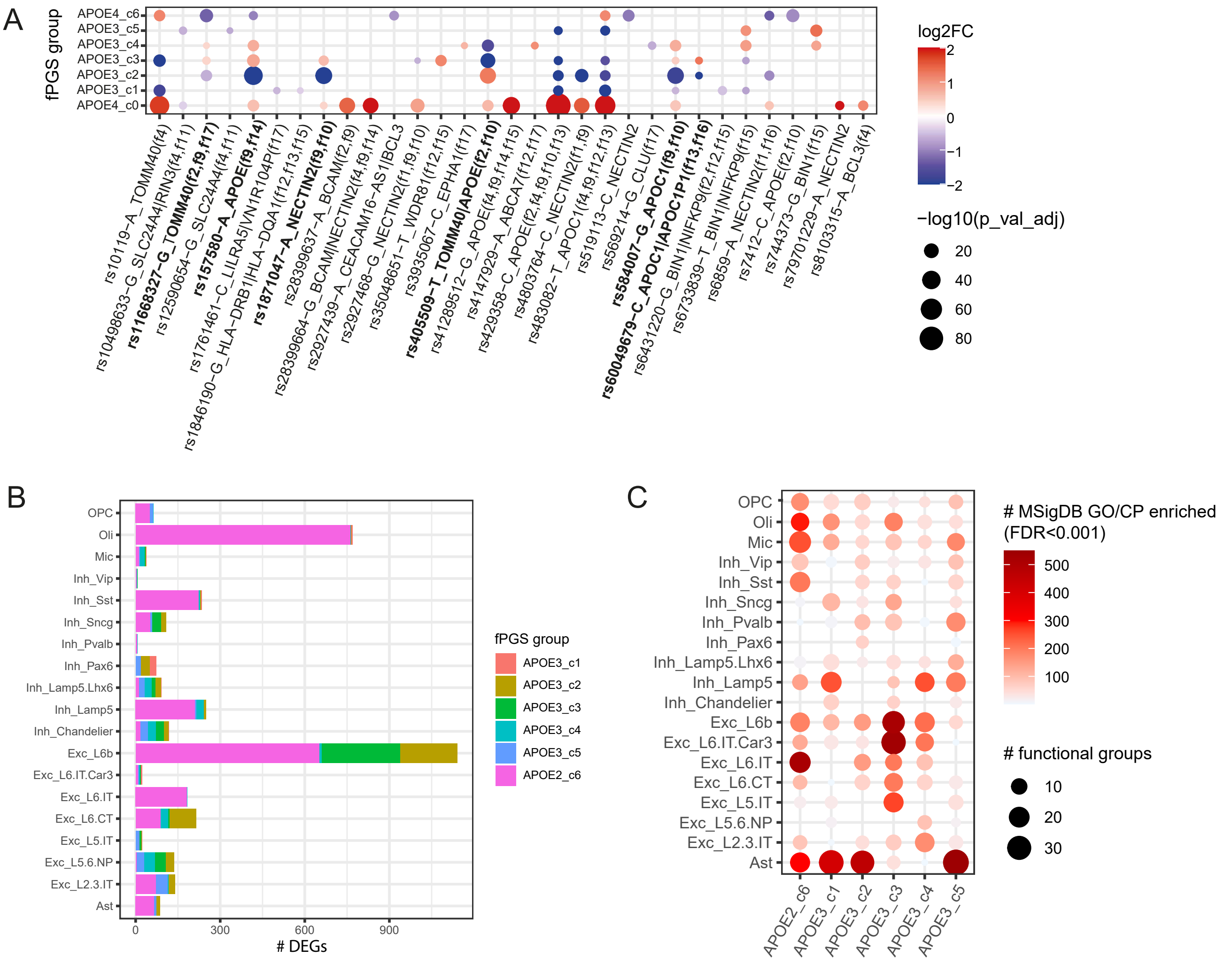

Supplementary Figure 5

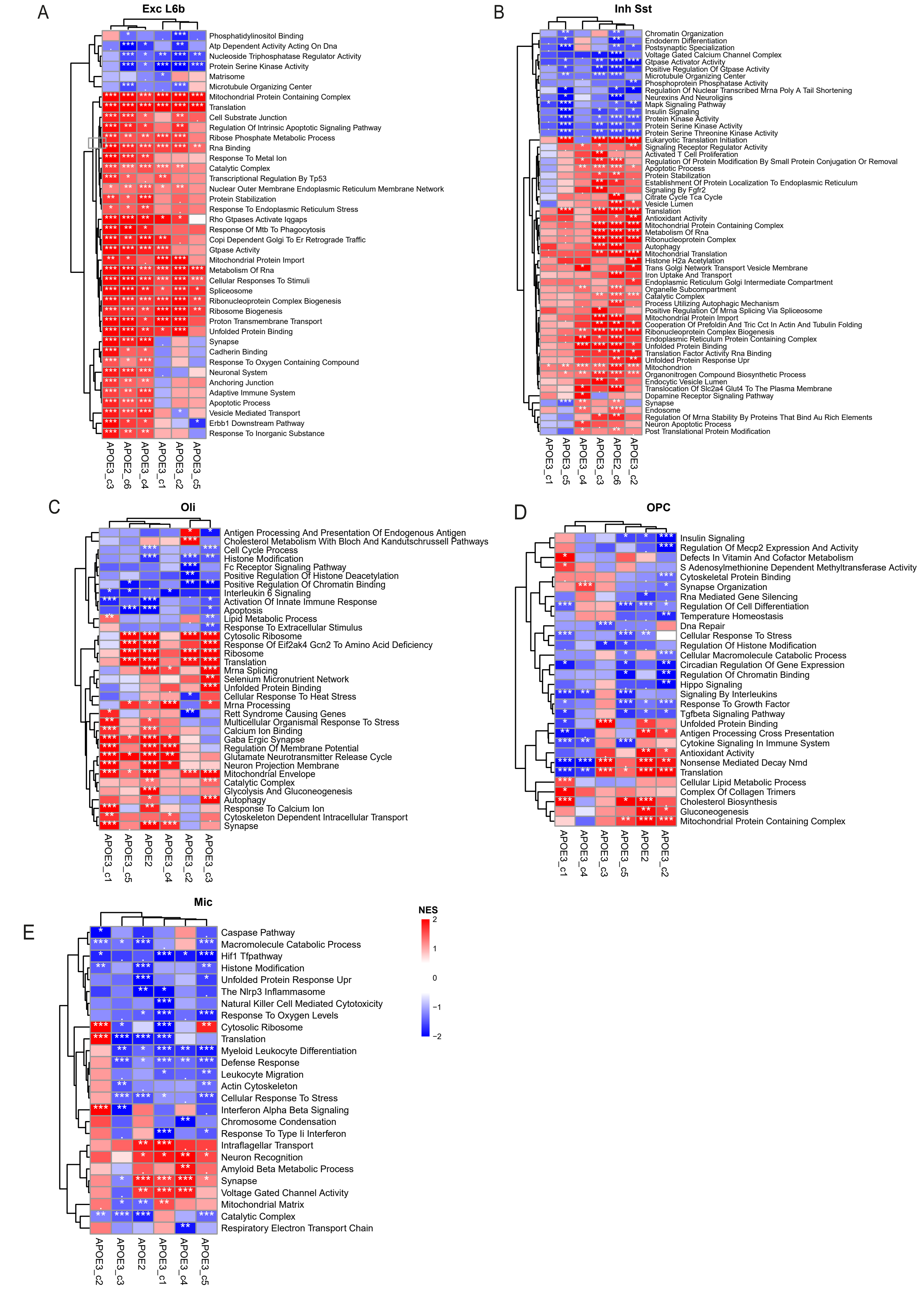
